# Mutant HTT protein decreases with CAG repeat expansion: implications for therapeutics and bioassays

**DOI:** 10.1101/2024.06.11.598410

**Authors:** Christian Landles, Georgina F. Osborne, Jemima Phillips, Maria Canibano Pico, Iulia M. Nita, Nadira Ali, Jonathan R. Greene, Kirupa Sathasivam, Gillian P. Bates

**Affiliations:** Department of Neurodegenerative Disease and Huntington’s Disease Centre, Queen Square Institute of Neurology, University College London, Queen Square, London WC1N 3BG, UK; Rancho BioSciences, San Diego, California 92127, USA

**Author notes:** Correspondence to: Gillian Bates and Christian Landles, Department of Neurodegenerative Disease and Huntington’s Disease Centre, Queen Square Institute of Neurology, UCL, Queen Square, London, WC1N 3BG, UK.

**Keywords:** Huntington’s disease, mutant huntingtin protein, huntingtin bioassay, HTT1a protein, knock-in mouse models of Huntington’s disease

## Abstract

Huntington’s disease is an inherited neurodegenerative disorder caused by a CAG repeat expansion that encodes a polyglutamine tract in the HTT protein. The mutant CAG repeat is unstable and expands in specific brain cells and peripheral tissues throughout life. Genes involved in the DNA mismatch repair pathways, known to act on expansion, have been identified as genetics modifiers, therefore, it is the rate of somatic CAG repeat expansion that drives the age of onset and rate of disease progression. In the context of an expanded CAG repeat, the *HTT* pre-mRNA can be alternatively processed to generate the *HTT1a* transcript, that encodes the aggregation prone and highly pathogenic HTT1a protein. This may be a mechanism through which somatic CAG repeat expansion exerts its pathogenic effects, as the longer the CAG repeat, the more *HTT1a* and HTT1a is produced.

The allelic series of knock-in mouse models: *Hdh*Q20, *Hdh*Q50, *Hdh*Q80, *Hdh*Q111, CAG140 and zQ175 with polyQ expansions of 20, 50, 80, 111 140 and ∼190 can be used to model the molecular and cellular consequences of CAG repeat expansion within a single neuron. By western blot of cortical lysates, we found that mutant HTT levels decreased with increasing CAG repeat length; mutant HTT was only 23% and 10% of wild-type levels in CAG140 and zQ175 cortices, respectively. To identify the optimal bioassays for detecting the full-length HTT and HTT1a isoforms, we interrogated the pairwise combinations of seven well-characterized antibodies on both the HTRF and MSD platforms. In total we tested 32 HTRF and 32 MSD assays to detect ‘full-length mutant HTT’, HTT1a, ‘total mutant HTT’ (full-length HTT and HTT1a) and ‘total full-length HTT’ (mutant and wild type). None of these assays recapitulated the full-length mutant HTT levels as measured by western blot. We recommend using isoform- and species-specific assays that detect either full-length mutant HTT, HTT1a or wild-type HTT as opposed to those that detect more than one isoform simultaneously.

Our finding that as the CAG repeat expands, full-length mutant HTT levels decrease, whilst *HTT1a* and HTT1a levels increase has implications for therapeutic strategies. If mutant HTT levels in cells containing (CAG)_200_ are only 10% of wild-type, HTT-lowering strategies targeting full-length *HTT* at sequences 3’ to intron 1 *HTT* will predominantly lower wild-type HTT, as mutant HTT levels in these cells are already depleted. These data support a therapeutic strategy that lowers *HTT1a* and depletes levels of the HTT1a protein.

## INTRODUCTION

Huntington’s disease is an inherited neurodegenerative disorder that manifests with motor, cognitive and psychiatric impairments.^1^ It is caused by an expanded CAG repeat in exon 1 of the huntingtin gene (*HTT*) that results in an expanded glutamine tract in the huntingtin protein (HTT).^2^ Repeats of (CAG)_40_ or more, as measured in blood, are fully penetrant, individuals with (CAG)_35_ or less remain unaffected,^3^ whereas, individuals with expansions of around (CAG)_65_ or more develop symptoms in childhood or adolescence.^4^ The CAG repeat is somatically unstable, expanding with age in specific brain regions.^5–7^ Given that several DNA mismatch repair pathway genes, known to modify somatic CAG repeat expansion, have been identified as genetic modifiers of Huntington’s disease, this phenomenon is considered to drive the age of onset and rate of disease progression.^8–11^ *HTT* pre-mRNAs with expanded CAG repeats, can be alternatively processed to generate the *HTT1a* transcript that encodes the aggregation-prone and highly pathogenic HTT1a protein.^12, 13^ The levels of *HTT1a* increase with increasing CAG repeat length and therefore, may be a mechanism through which somatic CAG expansion exerts its pathogenic effects.

It is essential that the levels of HTT protein isoforms can be measured in disease models and clinical samples to allow the interpretation of mechanistic studies and preclinical and clinical interventions. This is particularly important for potential therapies that target *HTT* directly through *HTT* lowering approaches.^14^ Assays that detect soluble or aggregated HTT isoforms have been developed by utilizing pairs of anti-HTT antibodies. For cell lysates or tissue extracts from Huntington’s disease models, assays have been developed using homogeneous time resolved fluorescence (HTRF) (also termed TR-FRET),^15, 16^ Mesoscale Discovery (MSD)^17, 18^ and amplified luminescent proximity homogeneous assay (AlphaLISA)^19^ platforms. More sensitive single molecule counting (SMC) assays have been needed to detect HTT in CSF.^20, 21^

In this study, we investigated the effect of the Huntington’s disease mutation on mutant HTT protein levels as well as the comparative performance of HTRF and MSD assays designed to detect soluble HTT isoforms. We used cortical lysates from heterozygous and homozygous *Hdh*Q20, *Hdh*Q50, *Hdh*Q80, *Hdh*Q111, CAG140 and zQ175 knock-in mice, together with their wild-type littermates at 11 weeks of age and included lysate from YAC128 transgenic mice, to control for the detection of HTT levels above wild-type. This allelic series of knock-in mice can be used to model changes in mutant HTT levels in response to large somatic CAG expansions within a single neuron.

We used western blotting to estimate full-length wild-type and mutant HTT levels in cortical lysates, and then assessed the ability of 12 HTRF and 12 MSD assays to measure ‘full-length mutant HTT’, two HTRF and two MSD assays to measure HTT1a, six HTRF and six MSD assays to measure ‘total mutant HTT’ (full-length and HTT1a) and 12 HTRF and 12 MSD assays to measure ‘total full-length HTT’ (mutant and wild type). The western blots demonstrated that mutant HTT levels in lysates from homozygous *Hdh*Q20 and *Hdh*Q50 mice were equivalent to wild type. However, as the CAG repeat in the knock-in alleles of the *Hdh*Q80, *Hdh*Q111, CAG140 and zQ175 lines increased, full-length mutant HTT levels in the homozygous knock-in mice decreased to approximately 10% of wild type in zQ175 cortices. In contrast, HTT1a protein levels increased. This finding has significant implications for HTT lowering therapeutic interventions and our data stress that it is essential to understand the effect of polyQ length on the performance of HTT bioassays before using them for mechanistic or therapeutic applications in the preclinical or clinical setting.

## MATERIALS AND METHODS

### Ethics Statement

All procedures were performed in accordance with the Animals (Scientific Procedures) Act, 1986, and approved by the University College London Ethical Review Process Committee.

### Mouse Breeding and Maintenance

Heterozygous, homozygous and wild-type littermates were imported from the CHDI Foundation colonies at the Jackson Laboratory (Bar Harbor, Maine) for the following knock-in lines on a C57BL/6J background: *Hdh*Q20, *Hdh*Q50, *Hdh*Q80 and *Hdh*Q111,^22^ CAG140^23^ and zQ175^24, 25^ and housed at UCL until 11 weeks of age. YAC128 mice^26^ were bred at UCL by pairing YAC128 males with C57BL/6J females (Charles River, Netherlands) and tissues collected at 9 weeks of age. Mouse husbandry and health monitoring were as previously described.^27^ Animals were kept in individually ventilated cages with Aspen Chips 4 Premium bedding (Datesand) and environmental enrichment comprising chew sticks and a play tunnel (Datesand). All mice had constant access to water and food (Teklad global 18% protein diet, Envigo, the Netherlands). Temperature was regulated at 21°C ± 1°C and mice were kept on a 12 h light/dark cycle. The facility was barrier-maintained and quarterly non-sacrificial FELASA (Federation of European Laboratory Animal Science Associations) screens found no evidence of pathogens. Tissues were harvested, snap frozen in liquid nitrogen and stored at -80°C.

### Genotyping and CAG repeat sizing

YAC128 mice were genotyped as previously described.^28^ The polyQ repeat of 125 glutamines in YAC128 mice is encoded by (CAG)_23_(CAA)_3_CAGCAA(CAG)_80_(CAA)_3_CAGCAA(CAG)_10_CAACAG which is stable on germline transmission.^29^ For imported mice, tail biopsies were collected upon sacrifice for genotype confirmation. The *Hdh*Q20, *Hdh*Q50, *Hdh*Q80 and *Hdh*Q111 mice were genotyped as previously described for *Hdh*Q20^28^ and the CAG140 and zQ175 mice as described for zQ175.^28^ CAG repeat sizing had been performed by the CHDI Foundation before shipment to UCL and CAG lengths are summarized in **Supplementary Table 1**.

### Antibodies

Antibodies are summarized in **Supplementary Table 2**, and their epitope locations on the HTT protein are shown in **Fig. 1A**.

**Figure 1.**
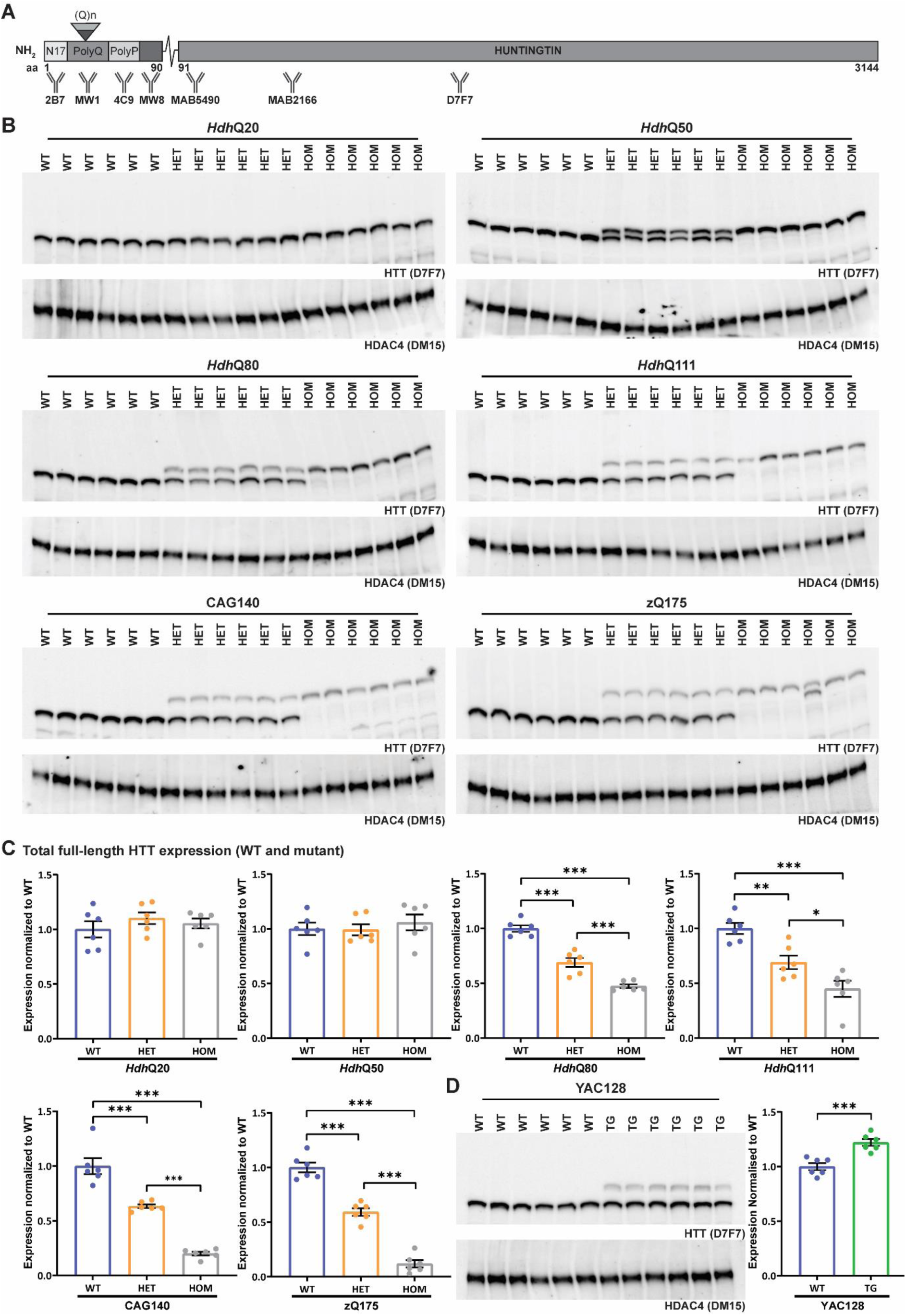
Full-length mutant HTT levels decrease with increasing CAG repeat length. **(A)** Schematic showing the epitope locations of the HTT antibodies used in this study. Details of immunogens and antibody epitopes are given in **Supplementary Table 2**. **(B)** Western blots of full-length HTT, as detected by D7F7 in cortical lysates from wild-type, heterozygous, and homozygous mice (n = 6 / genotype) for each of the knock-in mouse lines: *Hdh*Q20, *Hdh*Q50, *Hdh*Q80, *Hdh*Q111, CAG140 and zQ175. The DM15 antibody against HDAC4 was used as a loading control. The full-length blots and their quantification are shown in **Supplementary Figs. 4–9**. **(C)** Quantification of the levels of total full-length HTT (wild-type and mutant) from the western bots shown in (B). **(D)** Western blot of full-length HTT, as detected by D7F7 in cortical lysates from wild-type and YAC128 transgenic mice (n = 6 / genotype). DM15 antibody against HDAC4 was used as a loading control. The full-length blot and its quantification are shown in **Supplementary Fig. 10**. Statistical analysis was by Student’s *t*-test and one-way ANOVA with Bonferroni *post hoc* correction. Error bars are mean ± SEM. \**P* ≤ 0.05, \*\**P* ≤ 0.01, \*\*\**P* ≤ 0.001. aa = amino acid, polyQ = polyglutamine, polyP = polyproline, WT = wild type, HET = heterozygote, HOM = homozygote, TG = transgenic.

### Western Blotting

Cortical lysates were prepared in ice cold RIPA buffer (50 mM TRIS pH 8.0, 150 mM NaCl, 1% NP40, 0.5% Na-cholate, 0.1% SDS), NP40 buffer (50 mM TRIS pH 7.5, 150 mM NaCl, 1% NP40) or HEPES buffer (50 mM HEPES pH 7.0, 150 mM NaCl, 10 mM EDTA, 1% NP40, 0.5% Na-cholate, 0.1% SDS) with c0mplete protease inhibitors (Roche). Proteins were denatured in Laemmli loading buffer as previously described,^30^ 25 µg of protein was separated by 7.5% SDS-polyacrylamide acrylamide gel electrophoresis (SDS-PAGE) (Criterion, Bio-Rad) and blotted onto nitrocellulose membranes. Primary antibody dilution was 1:1000 for HTT (D7F7) and 1:1000 for HDAC4 (DM15), blots were immunoprobed in PBST (phosphate buffered saline, 0.1% TWEEN 20) or TBST (TRIS-buffered saline, 0.1% TWEEN 20), and detected by chemiluminescence, as described previously.^31^ Quantification of western blots was performed using the Image Lab software (Bio-Rad).

### HTT bioassay protein lysate preparation

A 10% (w/v) total protein cortical homogenate was prepared in ice-cold bioassay buffer (phosphate buffer saline (PBS), 1% Triton-X-100) as previously described (n = 6 / genotype).^28^ Lysates were snap frozen and used for assays the following day.

### Homogenous time resolved FRET (HTRF) assay

Cortical homogenates to a final volume of 10 μL were pipetted in triplicate into a 384-well (pure white, low volume, conical) proxiplate (Greiner Bio-One). Lysate dilutions and antibody concentrations are summarized in **Supplementary Table 3**. For HTRF assays antibodies were added per well in 5 μL HTRF detection buffer (50 mM NaH_2_PO_4_, 0.2 M KF, 0.1% bovine serum albumin (BSA), 0.05% Tween-20) with cOmplete protease inhibitor cocktail tablets (Roche). Plates were incubated for 3 h on an orbital shaker (250 rpm) at room temperature, before reading on an EnVision (Revvity) plate reader using optimized HTRF detection parameters described previously.^28^

### Meso Scale Discovery (MSD) conjugation assay

Cortical homogenates to a final volume of 18 μL were pipetted in triplicate into 384-well, 1-spot custom printed MSD plates. MSD plates were run exactly according to the manufacturer’s recommended conditions. Lysate dilutions are summarized in **Supplementary Table 4** and all incubations were performed at room temperature on a jitterbug shaker (Boekel Scientific) at 1,000 rpm. Wells were blocked by incubating overnight in 45 μL 3% BSA in PBS. HTT proteins were then captured for 3 h at the default MSD anti-huntingtin capture concentration of ∼2 mg/mL ± 15% per well and detected for 1.5 h using 2.5 μg/mL anti-huntingtin Sulfo-tag, except for MW1 Sulfo-tag which was used at 1.5 μg/mL. Between each step, wells were washed three times in PBS-Tween buffer (PBS, 0.05 % Tween-20). For detection, 45 μL MSD GOLD read buffer was added per well and read on an MSD plate reader (Meso Scale Discovery).

### RNA extraction and real time quantitative PCR

Cortical tissue was homogenized in Qiazol lysis reagent in lysing matrix tubes D (MP Biomedicals) three times for 30 s using the Fastprep-24 (MP Biomedicals). Total RNA was extracted, DNase I treated, and reverse transcribed with oligodT_18_ (Invitrogen) for *Htt1a* and random hexamers (Invitrogen) for full-length *Htt* as previously described.^32^

Primers and probes for the full-length *Htt* and *Htt1a* transcripts (**Supplementary Table 5**) and the custom Taqman assays for mouse reference genes (*Atp5b*, *Sdha*, *Eif4a2*) were from Thermo Fisher Scientific. Reactions were performed in triplicate. Each 12 μL reaction consisted of 3 μL of 1:10 diluted cDNA, TaqMan Fast Advanced Mix (Applied Biosystems) and primer probe mix distributed into 384-well thin-walled white PCR plates (Bio-Rad). Plates were sealed with Microseal ‘B’ seals (Bio-Rad), centrifuged at 800 x *g* for 30 s and amplified in a Bio-Rad CFX Opus 384 RealTtime PCR machine as follows: 40 s at 95 °C, 40 x (7 s at 95 °C, 20 s at 60 °C). C*q* values deviating by 0.5 from the mean were excluded from further analysis. Data for genes of interest were normalized to the geometric mean of the reference genes and the 2^-ΔCt2^ was calculated.^33^

### Statistical analysis

Data were screened for outliers using Grubb’s test and outliers were removed before between-group comparisons. All data sets were tested for a normal Gaussian distribution (Shapiro-Wilk test). Statistical analysis was performed using either Student’s *t*-test or one-way ANOVA with Bonferroni *post hoc* test. All statistical analyses were performed, and graphs prepared using GraphPad Prism (v10).

## RESULTS

We set out to determine the effect of increasing CAG repeat length on levels of mutant HTT, and to investigate how best to detect HTT protein isoforms by HTRF and MSD assays. All analyses were performed using fresh cortical lysates from homozygous, heterozygous, and wild-type mice from *Hdh*Q20, *Hdh*Q50, *Hdh*Q80, *Hdh*Q111, CAG140 and zQ175 knock-in mouse colonies at 11 weeks of age (in total, two sets of n = 3 mice / gender / genotype for all western, qPCR and bioassay data). All knock-in lines had been generated by replacing wild-type mouse exon 1 *Htt* with a mutant version of human exon 1 *HTT*.^22, 23^ Cortical lysates were also prepared from transgenic YAC128 mice and their wild-type littermates at 9 weeks of age (n = 6 / genotype), to control for the detection of mutant HTT in addition to endogenous wild-type levels.

The epitope locations for the anti-HTT antibodies are illustrated in **Fig. 1A**. The D7F7 antibody was used to detect full-length HTT levels on western blots. For HTRF and MSD assays, pairwise antibody combinations were selected that would be predicted to detect ‘full-length mutant HTT’ (MW1 or 4C9 with MAB5490, MAB2166 or D7F7); HTT1a (2B7 or MW1 with MW8), ‘total mutant HTT’ (HTT1a and full-length mutant HTT) (2B7, MW1 and 4C9) and ‘total full-length HTT’ (mutant and wild type) (2B7, MAB5490, MAB2166 and D7F7). Depending on the antibody location, the full-length HTT assays will also detect either all, or a subset of, proteolytic HTT fragments. For each antibody pair, the two antibodies were used in both donor and acceptor orientations for HTRF and in both capture and detection orientations for MSD.

### Mutant full-length HTT levels decrease with increasing CAG repeat length

For western blot experiments, we began by preparing cortical lysates in three different detergent-containing lysis buffers: RIPA, NP40 and HEPES from the same wild-type and zQ175 mice (n =2 / genotype). Blots were immunoprobed for HTT with D7F7 in either PBST or TBST. The signals were marginally stronger with less background for the RIPA lysates and for blots immunoprobed in PBST (**Supplementary Fig. 1**).

Cortical lysates were prepared in RIPA buffer from wild-type, heterozygous and homozygous mice for each of the knock-in colonies, separated by 7.5% SDS-PAGE and blots were probed with D7F7 for full-length HTT, and with DM15 for HDAC4 as a loading control (**Fig. 1B**). Expanded polyQ tracts retard HTT migration, and consequently, mutant HTT could be distinguished from wild type for all lines except for *Hdh*Q20 (**Fig. 1B**). Blots were quantified to compare total levels of HTT (wild-type and mutant HTT) in cortical lysates between wild-type, heterozygous and homozygous mice for each line (**Fig. 1C and Supplementary Fig. 2A**). The levels of mutant and wild-type HTT were equivalent in the *Hdh*Q20 and *Hdh*Q50 lines. In contrast, comparison of the wild-type and homozygous mutant signals indicated that the level of total HTT in the *Hdh*Q80, *Hdh*Q111, CAG140 and zQ175 cortical lysates had decreased with increasing CAG/polyQ length (**Fig. 1C and Supplementary Fig. 2A**). HTT signals for each knock-in line could be compared within a blot but not between blots. Therefore, to better demonstrate the reduction in mutant HTT between the *Hdh*Q111, CAG140 and zQ175 lines, homozygote lysates were separated on the same gel (**Supplementary Fig. 3**). The level of mutant HTT in the *Hdh*Q111, CAG140 and zQ175 lines was approximately 50%, 23% and 10% of wild-type HTT, respectively (**Fig. 3D and Supplementary Fig. 3C**).

**Figure 2.**
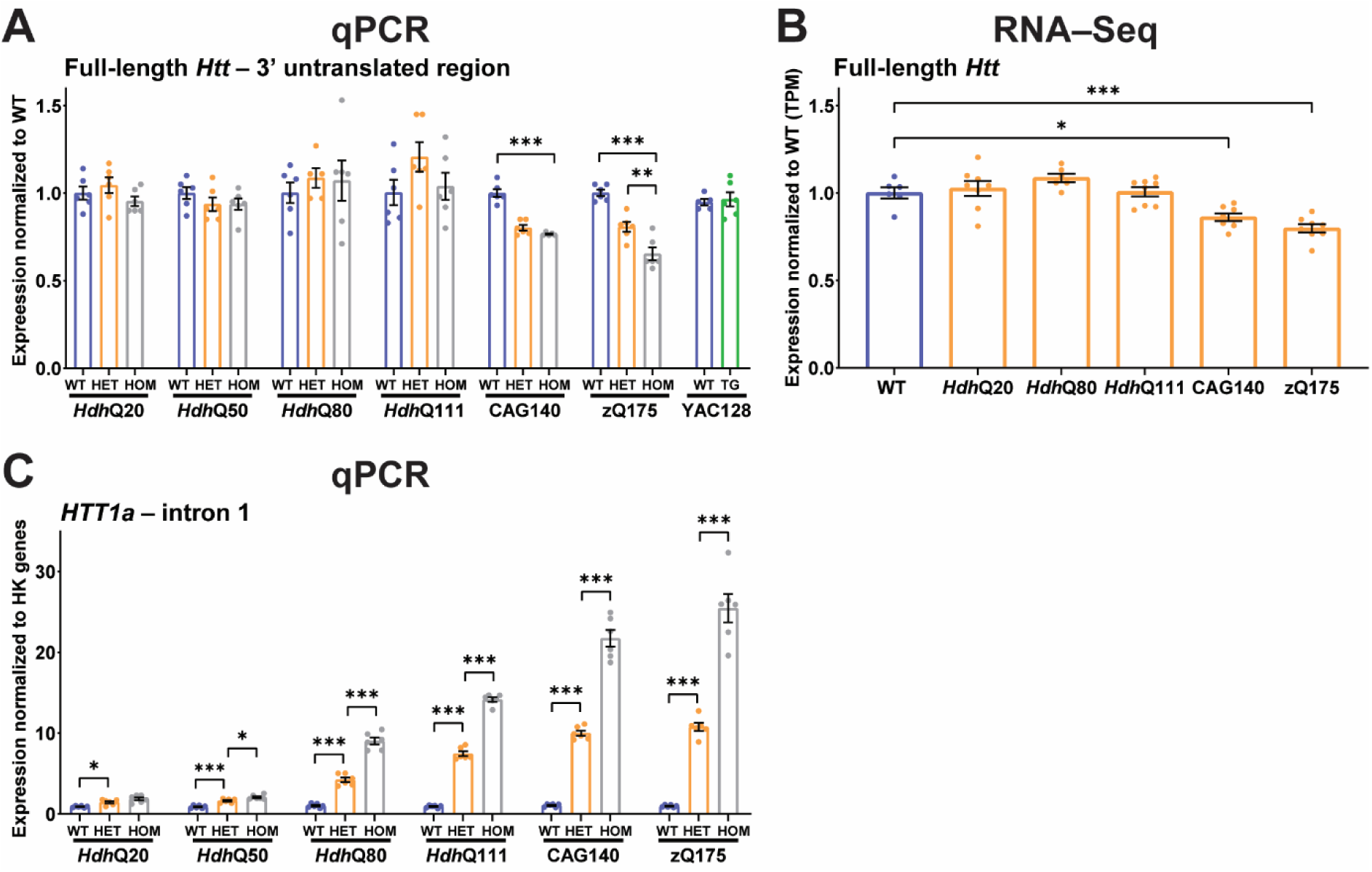
Changes in *Htt* transcript levels do not account for the reduction in mutant HTT levels in the knock-in mouse lines. **(A)** qPCR for full-length *Htt* transcript levels in cDNA prepared from the cortex of wild-type, heterozygous and homozygous knock-in mice and from YAC128 and wild-type mice. Cortical full-length *Htt* mRNA levels were comparable between wild-type, heterozygous and homozygous mice for the *Hdh*Q20, *Hdh*Q50, *Hdh*Q80 and *Hdh*Q111 lines. There was a reduction in full-length *Htt* in the heterozygous and homozygous CAG140 and zQ175 mice. This qPCR assay amplifies mouse *Htt* and therefore does not detect the human *HTT* transgene in YAC128 mice but demonstrates that wild-type *Htt* levels were not altered. **(B)** Determination of comparative full-length *Htt* levels in wildztype and heterozygous *Hdh*Q20, *Hdh*Q80, *Hdh*Q111, CAG140 and zQ175 cortex from RNA-seq datasets.^34^ There was no difference in the level of full-length *Htt* levels between wild-type and heterozygous *Hdh*Q20, *Hdh*Q80 and *Hdh*Q111 mice. *Htt* levels were reduced by approximately 14 % and 20% of wild-type levels in the CAG140 and zQ175 heterozygotes, respectively. **(C)** qPCR for *Htt1a* transcript levels in cDNA prepared from the cortex of wild-type, heterozygous and homozygous knock-in mice. The level of *Htt1a* increases with increasing CAG repeat length and homozygous levels are approximately twice that in heterozygotes. Statistical analysis was Student’s *t*-test or one-way ANOVA with Bonferroni *post hoc* correction, mean ± SEM. \**P* ≤ 0.5, \*\**P* ≤ 0.01, \*\*\**P* ≤ 0.001. WT = wild type, HET = heterozygote, HOM = homozygote, HK = housekeeping gene, TG = transgenic, TPM = transcripts per million.

**Figure 3.**
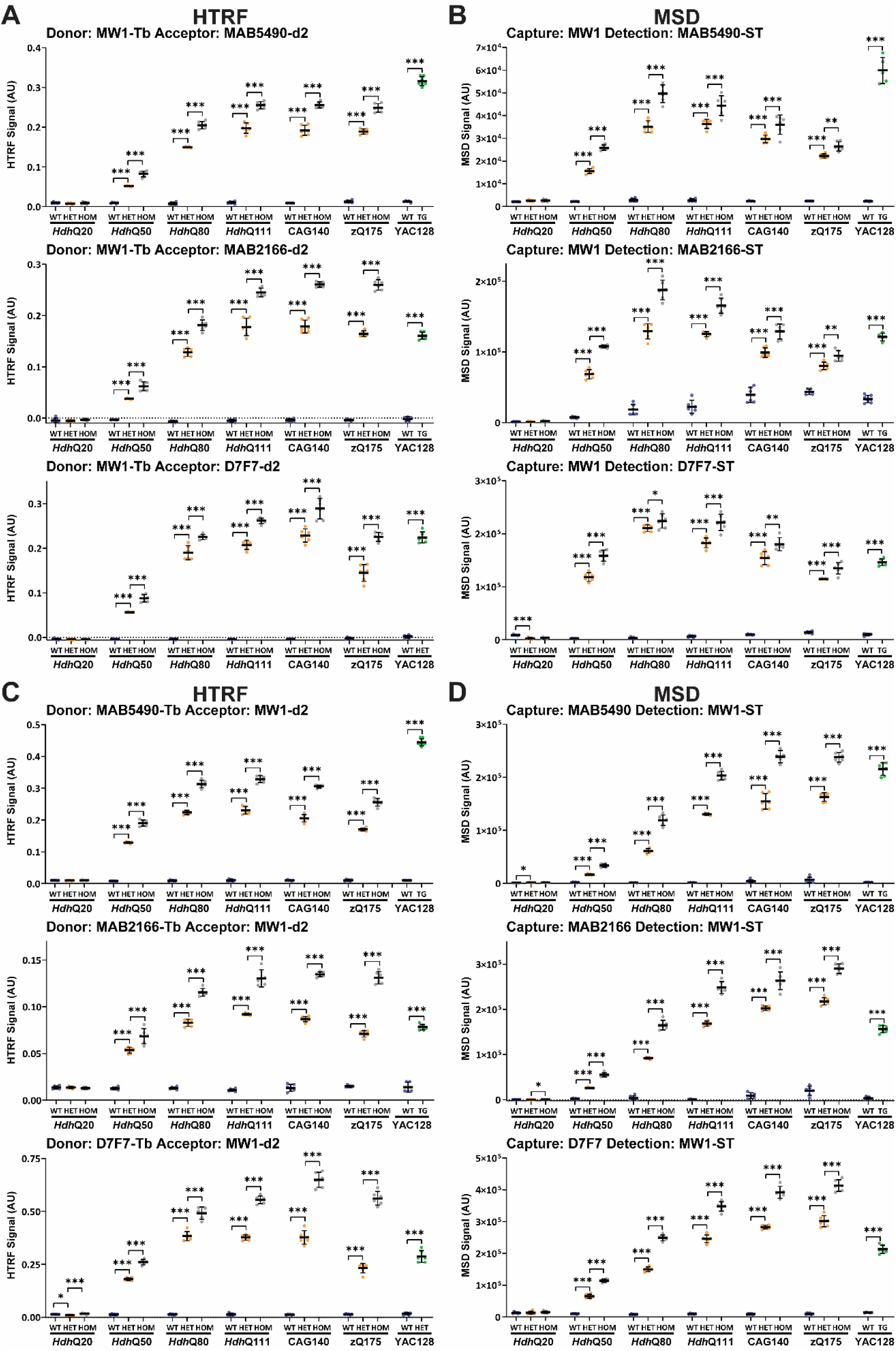
PolyQ-length dependence of HTRF and MSD assays that use MW1 and detect soluble full-length mutant HTT. Antibody pairings of MW1-MAB5490, MW1-MAB2166 and MW1-D7F7 were tested by HTRF **(A)** and MSD **(B)** and of MAB5490-MW1, MAB2166-MW1 and D7F7-MW1 were tested by HTRF **(C)** and MSD **(D)** using cortical lysates from wild-type, heterozygous and homozygous *Hdh*Q20, *Hdh*Q50, *Hdh*Q80, *Hdh*Q111, CAG140 and zQ175 mice at 11 weeks of age and YAC128 and wild-type littermates at 9 weeks of age (n = 6 / genotype). Statistical analysis was Student’s *t*-test or one-way ANOVA with Bonferroni *post hoc* correction per mouse line, mean ± SEM. \**P* ≤ 0.5, \*\**P* ≤ 0.01, \*\*\**P* ≤ 0.001. WT = wild type, HET = heterozygote, HOM = homozygote, TG = transgenic.

The HTT signal in YAC128 cortical lysates was approximately 120% of wild-type HTT; mutant HTT levels were approximately 40% of that arising from an endogenous wild-type allele (**Fig. 1D and Supplementary Fig. 2A and C**). Quantification of the wild-type HTT signals in heterozygous knock-in lysates confirmed that these were approximately 50% of that in wild-type mice and the level of wild-type HTT in YAC128 lysates was at endogenous levels (**Supplementary Fig. 2B**). Quantification of the mutant HTT signals in homozygous knock-in lysates confirmed that these were approximately double that detected in the heterozygous lysates for all lines (**Supplementary Fig. 2C**).

### The reduction in mutant HTT cannot be attributed to decreased *Htt* transcript levels

We prepared cortical cDNA from wild-type, heterozygous and homozygous mice from the knock-in lines and from YAC128 and wild-type littermates. Quantitative real-time PCR (qPCR) indicated that full-length *Htt* levels in heterozygous and homozygous *Hdh*Q20, *Hdh*Q50, *Hdh*Q80 and *Hdh*Q111 cortices were comparable to wild type but decreased in CAG140 and zQ175 (**Fig. 2A**). This reduction was confirmed by comparing full-length *Htt* reads in the previously published RNA-seq data^34^ from the allelic series of knock-in mice; mRNA levels being reduced by 14% and 20% of wild-type in heterozygous CAG140 and zQ175 cortex, respectively (**Fig. 2B**). In contrast, *Htt1a* levels increased with increasing CAG repeat length. The levels in homozygous mice were approximately double those in their heterozygous counterparts (**Fig. 2C**).

### Characterization of assays designed to detect ‘full-length mutant HTT’ (excluding the HTT1a protein)

The MW1 antibody recognizes an expanded polyQ tract and since it does not detect endogenous mouse HTT can be used to develop assays selective for mutant HTT.^15, 28^ Pairings between MW1 and antibodies C-terminal to HTT1a (MAB5490, MAB2166 and D7F7), will therefore detect ‘full-length mutant HTT’, as well as some proteolytic fragments dependent on which of these three antibodies has been chosen. We performed HTRF with MW1 as the donor antibody (**Fig. 3A**) and MSD with MW1 as the capture antibody (**Fig. 3B**) paired with MAB5490, MAB2166 or D7F7, as well as the reciprocal assays of HTRF with MW1 as the acceptor antibody (**Fig. 3C**) and MSD with MW1 as the detector antibody (**Fig. 3D**) paired with MAB5490, MAB2166 or D7F7. In all cases the assays were specific for mutant HTT (**Fig. 3**). Because MW1 binds to the polyQ tract, the assay signal might be expected to increase with increasing polyQ length. This occurred for all assays for polyQ sizes from Q20 to Q80, but from Q80 or Q111 onwards the signals either continued to increase, leveled out or decreased (**Fig.3**). In all cases, the signal was greater in homozygous as compared to heterozygous lysates. Therefore, the failure of the signal to continue to increase in heterozygotes with increasing polyQ length was not due to limiting antibody concentrations (**Fig. 3**). It was only in the *Hdh*Q50 cortex that, for some assays, the signal in homozygotes was double that obtained for the heterozygotes. The pattern by which the signal changed in response to increasing polyQ length was relatively comparable between HTRF and MSD assays, except for MW1-MAB2166 and MAB5490-MW1. The different assays gave very variable and inconsistent signals for the level of ‘full-length mutant HTT’ in the YAC128 cortex as compared to the knock-in lines, with assays utilizing MAB5490 giving the greatest signal (**Fig. 3**). In summary, the non-linear relationship between the length of the polyQ repeat and MW1 antibody binding may reflect conformational changes in very long polyQ tracts. These assays cannot be used to compare ‘full-length mutant HTT’ levels in tissues or biofluids containing HTT with variable polyQ lengths.

The 4C9 antibody is specific for the human polyproline-rich region in *HTT* exon 1 and can also be used for mutant HTT specific assays as the knock-in lines have a humanized exon 1.^16, 28^ To develop ‘full-length mutant HTT’ assays, we performed HTRF with 4C9 as the donor antibody (**Fig. 4A**) and MSD with 4C9 as the capture antibody (**Fig. 4B**) paired with MAB5490, MAB2166 or D7F7 as well as the reciprocal assays of HTRF with 4C9 as the acceptor antibody (**Fig. 4C**) and MSD with 4C9 as the detector antibody (**Fig. 4D**) paired with MAB5490, MAB2166 or D7F7. In all cases the assays were specific for mutant HTT (**Fig. 4**). The pattern by which the signal changed in response to increasing polyQ length was relatively comparable between HTRF and MSD assays for any antibody pairing. The heterozygous and homozygous signals decreased with increasing polyQ tract length for all assays, albeit to differing extents. The decrease in the heterozygote signal could not be due to limiting antibody concentrations as a greater signal was obtained for the homozygotes, and for many of the assays, the signal in the homozygote cortices was approximately double that in the heterozygotes (**Fig. 4**). Western blot analysis had indicated that full-length mutant HTT levels were the same as wild type in *Hdh*Q20 and *Hdh*Q50 cortex but decreased with increasing CAG repeat length from *Hdh*Q80 onwards (**Fig. 1C**). This change in mutant HTT levels was not precisely reflected by any of these assays. In some cases, the level of mutant HTT increased from *Hdh*Q20 to *Hdh*Q50, and in many cases, there was no difference between *Hdh*Q50 and *Hdh*Q80. Once again, different assays gave very variable signals for the level of full-length mutant HTT in the YAC128 cortex as compared to the knock-in lines, with assays utilizing MAB5490 giving the greatest signal (**Fig. 4**). The length of the polyQ repeat may influence the binding of the 4C9 antibody, because of the proximity of polyQ repeat and the 4C9 epitope.

**Figure 4.**
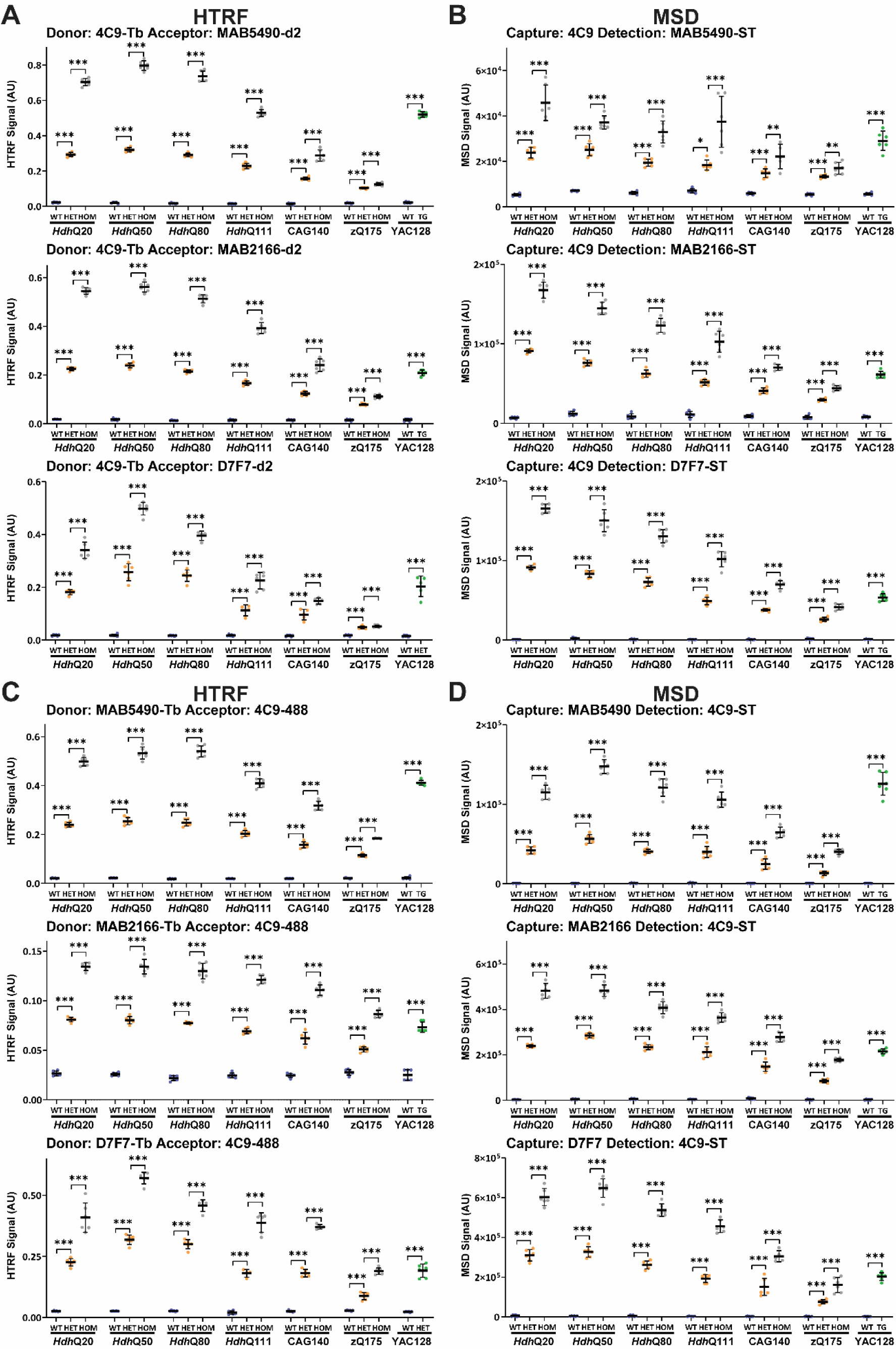
PolyQ-length dependence of HTRF and MSD assays that use 4C9 and detect soluble full-length mutant HTT. Antibody pairings of 4C9-MAB5490, 4C9-MAB2166 and 4C9-D7F7 were tested by HTRF **(A)** and MSD **(B)** and of MAB5490-4C9, MAB2166-4C9 and D7F7-4C9 were tested by HTRF **(C)** and MSD **(D)** using cortical lysates from wild-type, heterozygous and homozygous *Hdh*Q20, *Hdh*Q50, *Hdh*Q80, *Hdh*Q111, CAG140 and zQ175 mice at 11 weeks of age and YAC128 and wild-type littermates at 9 weeks of age (n = 6 / genotype). Statistical analysis was Student’s *t*-test or one-way ANOVA with Bonferroni *post hoc* correction per mouse line, mean ± SEM. \**P* ≤ 0.5, \*\**P* ≤ 0.01, \*\*\**P* ≤ 0.001. WT = wild type, HET = heterozygote, HOM = homozygote, TG = transgenic.

### Assays that detect the HTT1a protein (soluble or aggregated)

We have previously shown that MW8 is a neo-epitope antibody for the C-terminus of HTT1a under western blotting conditions^30^ and can be used to generate HTT1a specific HTRF, MSD and AlphaLISA assays^28^. To compare the effect of polyQ length on the detection of soluble HTT1a, we performed HTRF with 2B7 as the donor antibody (**Fig. 5A**) and MSD with 2B7 as the capture antibody (**Fig. 5B**) paired with MW8, as well as HTRF with MW1 as the donor antibody (**Fig. 5C**) and MSD with MW1 as the capture antibody (**Fig. 5D**) paired with MW8. For HTRF, the 2B7-MW8 assay (**Fig. 5A**) was much more sensitive than MW1-MW8 (**Fig. 5C**). For MSD, the 2B7-MW8 assay was very weak and not viable (**Fig. 5B**), with MW1-MW8 providing a better alternative (**Fig. 5D**). HTT1a levels increase with increasing CAG repeat (**Fig. 5A-D**) as would be predicted by the increasing levels of the *Htt1a* transcript (**Fig. 2C**) from which it is translated. The level of soluble HTT1a in the YAC128 cortex was comparable to that in an *Hdh*Q80 heterozygous mouse (**Fig. 5A-D**).

**Figure 5.**
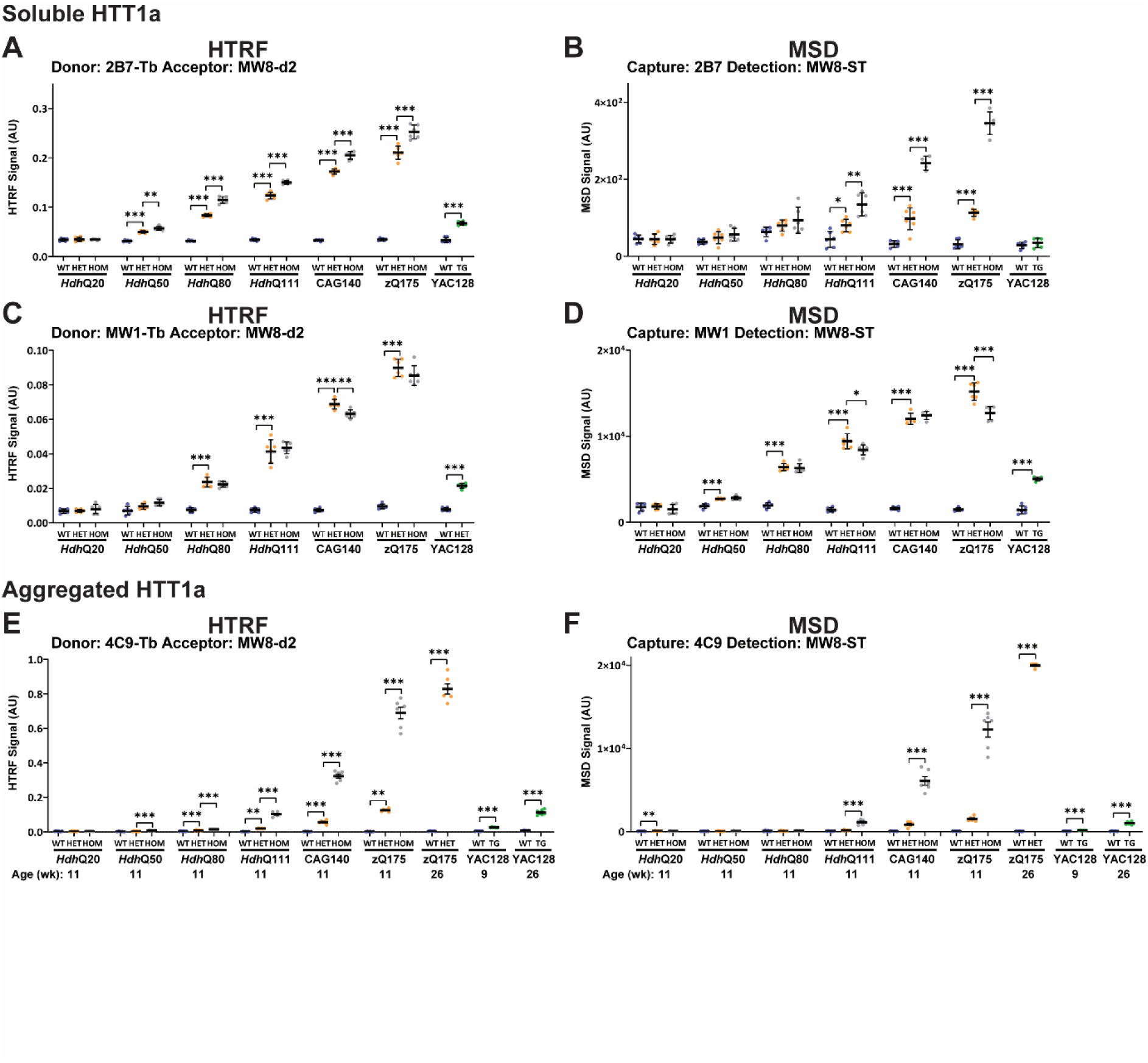
PolyQ-length dependence of HTRF and MSD assays that detect soluble and aggregated HTT1a. For soluble HTT1a, 2B7-MW8 was tested by HTRF **(A)** and MSD **(B)**, and MW1-MW8 was tested by HTRF **(C)** and MSD **(D)** using cortical lysates from wild-type, heterozygous and homozygous *Hdh*Q20, *Hdh*Q50, *Hdh*Q80, *Hdh*Q111, CAG140 and zQ175 mice at 11 weeks of age and YAC128 and wild-type littermates at 9 weeks of age (n = 6 / genotype). For aggregated HTT1a, 4C9-MW8 was tested by HTRF **(E)** and MSD **(F)** using cortical lysates from wild-type, heterozygous and homozygous *Hdh*Q20, *Hdh*Q50, *Hdh*Q80, *Hdh*Q111, CAG140 and zQ175 mice at 11 weeks of age, wild-type and zQ175 heterozygous mice at 26 weeks of age and wild-type and YAC128 mice at 9 and 26 weeks of age (n = 6 / genotype). Statistical analysis was Student’s *t*-test or one-way ANOVA with Bonferroni *post hoc* correction per mouse line, mean ± SEM. \**P* ≤ 0.5, \*\**P* ≤ 0.01, \*\*\**P* ≤ 0.001. WT = wild type, HET = heterozygote, HOM = homozygote, TG = transgenic.

We have previously shown that the 4C9-MW8 HTRF assay is specific for aggregated HTT1a.^35^ To investigate the comparable level of aggregated HTT1a we ran the 4C9-MW8 HTRF and MSD assays on lysates from the same cortices used for the soluble HTT1a assays (**Fig. 5E and F**). The HTRF assay was more sensitive than the MSD assay. At 11 weeks of age, statistically significant levels of HTT1a aggregation could be detected in heterozygous *Hdh*Q80 mice, and homozygous *Hdh*Q50 mice by HTRF, with the level in homozygous mice increasing more rapidly with polyQ length than that in heterozygous mice (**Fig. 5E and F**). Therefore, we would expect that the aggregation of HTT1a to have influenced the level of soluble HTT1a detected in **Fig. 5A-D** and may explain why the soluble HTT1a levels in homozygotes lysates were relatively comparable to those in heterozygotes lysates (**Fig. 5A-D**). The level of aggregation was similar in YAC128 cortex at 26 weeks of age to the cortex of heterozygous zQ175 mice at 11 weeks (**Fig. 5E and F**).

### Characterization of assays designed to detect ‘total mutant HTT’ (full-length HTT and HTT1a)

Total mutant HTT, comprising HTT1a, full-length mutant HTT and all proteolytic fragments can be detected by pairing two antibodies that both recognize epitopes within exon 1 of HTT. To investigate the effect of polyQ length on these assays, we performed HTRF with 2B7 as the donor antibody (**Fig. 6A**) and MSD with 2B7 as the capture antibody (**Fig. 6B**) paired with MW1 or 4C9; HTRF with MW1 as the donor antibody (**Fig. 6C**) and MSD with MW1 as the capture antibody (**Fig. 6D**) paired with 2B7 or 4C9; HTRF with 4C9 as the donor antibody (**Fig. 6E**) and MSD with 4C9 as the capture antibody (**Fig. 6F**) paired with 2B7 or MW1. In all cases, the assays were specific for mutant HTT (**Fig. 6**). The signal was predominantly greater in cortical lysates from homozygous as compared to heterozygous mice, indicating that the reductions in the heterozygous signals with CAG repeat length could not have resulted from limiting antibody concentrations. To ensure that this was also the case for the MW1-2B7 MSD assay in the *Hdh*Q111, CAG140 and zQ175 lysates (**Fig. 6D**), this assay was run through a series of dilutions for zQ175 lysates, and the homozygous signals were found to be greater than those for the heterozygotes (**Supplementary Fig. 11**).

**Figure 6.**
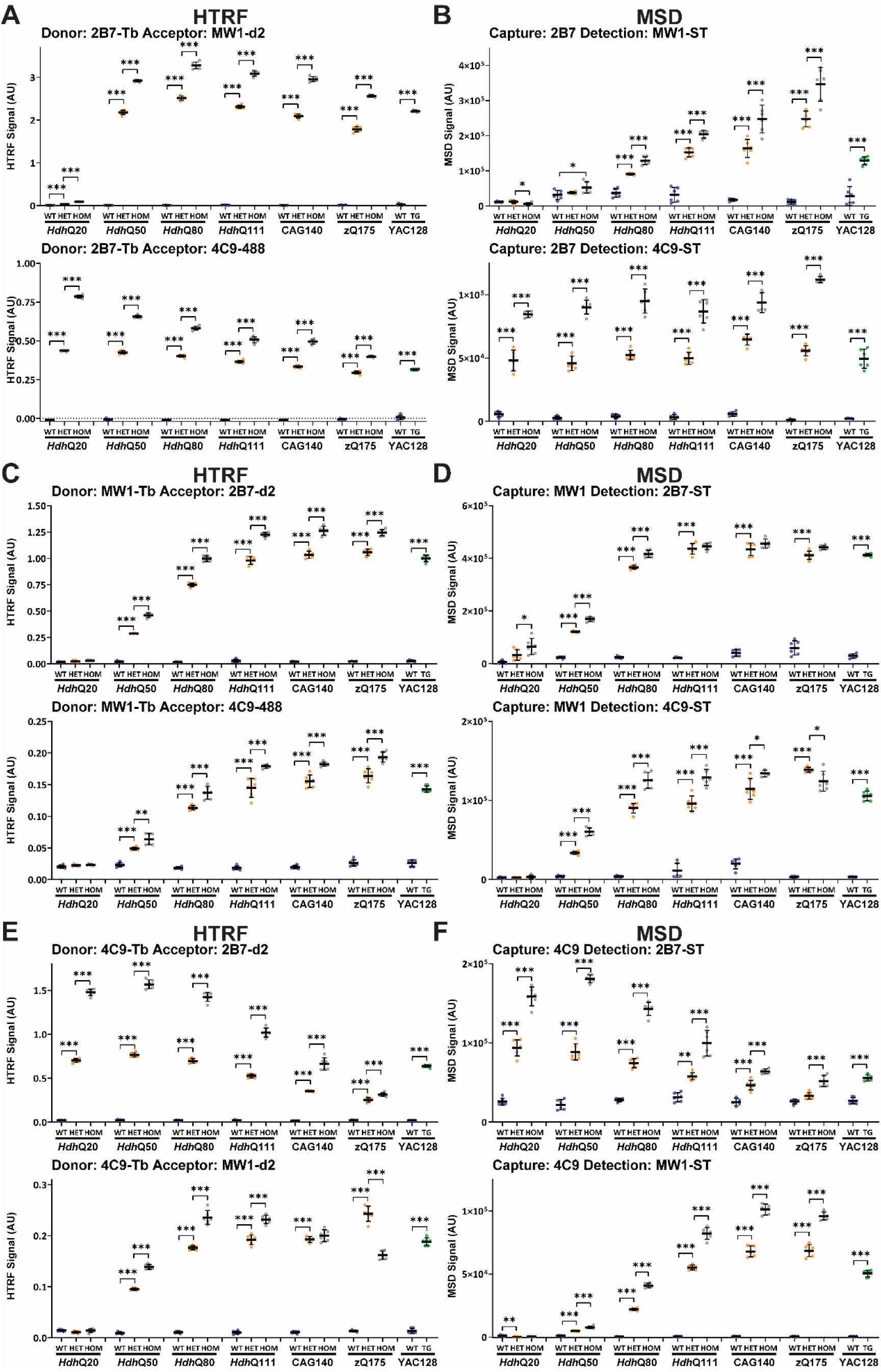
PolyQ-length dependence of HTRF and MSD assays that detect total soluble mutant HTT (full-length HTT and HTT1a). Antibody pairings of 2B7-MW1 and 2B7-4C9 were tested by HTRF **(A)** and MSD **(B)**, of MW1-2B7 and MW1-4C9 was tested by HTRF **(C)** and MSD **(D)**, and of 4C9-2B7 and 4C9-MW1 tested by HTRF **(E)** and MSD **(F)** using cortical lysates from wild-type, heterozygous and homozygous *Hdh*Q20, *Hdh*Q50, *Hdh*Q80, *Hdh*Q111, CAG140 and zQ175 mice at 11 weeks of age and YAC128 and wild-type littermates at 9 weeks of age (n = 6 / genotype). Statistical analysis was Student’s *t*-test or one-way ANOVA with Bonferroni *post hoc* correction per mouse line, mean ± SEM. \**P* ≤ 0.5, \*\**P* ≤ 0.01, \*\*\**P* ≤ 0.001. WT = wild type, HET = heterozygote, HOM = homozygote, TG = transgenic.

Comparison of the relative signals for these ‘total mutant HTT’ assays with the patterns obtained for ‘full-length mutant HTT’ (**Figs. 3 and 4**) and for HTT1a (**Fig. 5**) indicated that specific ‘total mutant HTT’ assays exhibited a preference for one of these isoforms. The most unexpected result was for the frequently used MSD assay, 2B7-MW1, which produced a pattern that tracked with HTT1a. The four HTRF and three other MSD assays utilizing MW1 described a pattern reminiscent of the ‘full-length mutant HTT’ assays (**Fig. 3**). The 4C9-2B7 HTRF and MSD assays also appear to track ‘full-length mutant HTT’, whereas the distinctive patterns of the 2B7-4C9 MSD and HTRF assays suggested that they are detecting both ‘full-length mutant HTT’ and HTT1a.

### Characterization of assays designed to detect ‘total full-length HTT’ (mutant and wild type)

Total full-length HTT (mutant and wild-type) can be detected by pairing 2B7 with antibodies that are located C-terminal to exon 1 HTT. We performed HTRF with 2B7 as the donor antibody (**Fig. 7A**) and MSD with 2B7 as the capture antibody (**Fig. 7B**) paired with MAB5490, MAB2166 or D7F7, as well as the reciprocal assays of HTRF with 2B7 as the acceptor antibody (**Fig. 7C**) and 2B7 as the detector antibody (**Fig. 7D**) paired with MAB5490, MAB2166 or D7F7. All antibody pairings showed much more inter-sample variability in the MSD data than the HTRF data (**Fig. 7**). For the HTRF assays, the wild-type signals for endogenous HTT were relatively comparable between mouse lines (**Fig. 7**).

**Figure 7.**
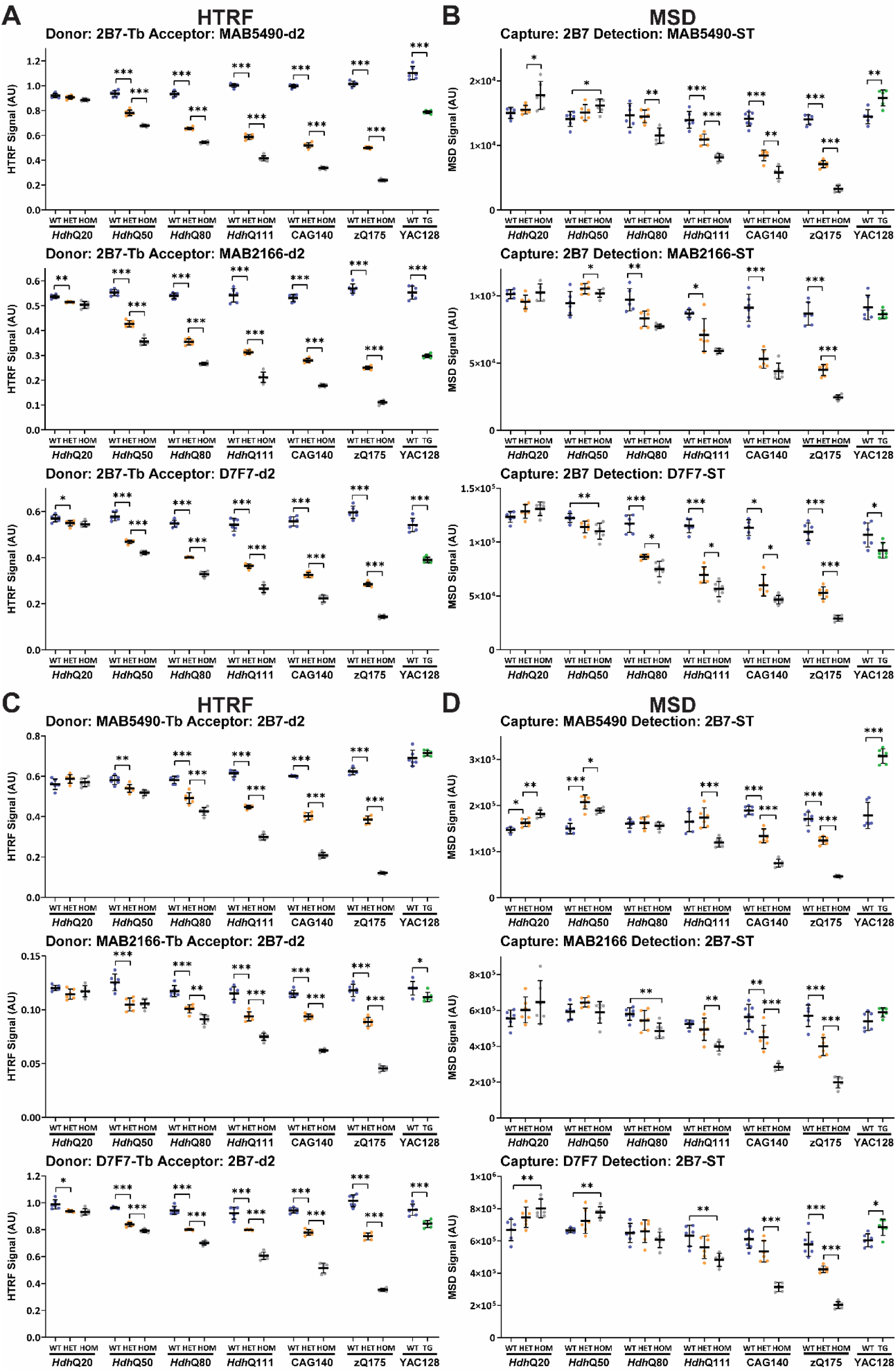
PolyQ-length dependence of HTRF and MSD assays that use 2B7 and detect total full-length HTT (mutant and wild-type). Antibody pairings of 2B7-MAB5490, 2B7-MAB2166 and 2B7-D7F7 were tested by HTRF **(A)** and MSD **(B)** and of MAB5490-2B7, MAB2166-2B7 and D7F7-2B7 were tested by HTRF **(C)** and MSD **(D)** using cortical lysates from wild-type, heterozygous and homozygous *Hdh*Q20, *Hdh*Q50, *Hdh*Q80, *Hdh*Q111, CAG140 and zQ175 mice at 11 weeks of age and YAC128 and wild-type littermates at 9 weeks of age (n = 6 / genotype). Statistical analysis was Student’s *t*-test or one-way ANOVA with Bonferroni *post hoc* correction per mouse line, mean ± SEM. \**P* ≤ 0.5, \*\**P* ≤ 0.01, \*\*\**P* ≤ 0.001. WT = wild type, HET = heterozygote, HOM = homozygote, TG = transgenic.

In general, the heterozygous and homozygous signals decreased with polyQ length, with the homozygote signals lower than the heterozygote, consistent with western blot data (**Fig. 1**). However, all HTRF assays indicated that the heterozygous and homozygous *Hdh*Q50 lysates contained less full-length HTT than wild type, which was inconsistent. Also, none of the HTRF assays registered that the YAC128 lysate contained more full-length HTT than wild type, and the three HTRF assays with 2B7 as donor, measured less HTT in YAC128 lysate than in wild type, which cannot be the case. An increase in total HTT in the YAC128 lysates was only apparent in the signals generated by the 2B7-MAB5490, MAB5490-2B7 and D7F7-2B7 MSD assays, but these assays also detected more full-length HTT in heterozygous and/or homozygous *Hdh*20 lysates as compared to wild type, which is again inconsistent with the western blot data. To investigate this further, the HTRF assays were repeated with 5% and 2.5% lysates. This demonstrated that the relative signals within and between lines were sensitive to lysate concentration (**Supplementary Fig. 12**), in a manner that is difficult to interpret.

Total full-length HTT can also be detected by pairing two antibodies that both recognize epitopes located C-terminal to exon 1 HTT. We performed HTRF with MAB5490 as the donor antibody (**Fig. 8A**) and MSD with MAB5490 as the capture antibody (**Fig. 8B**) paired with MAB2166 or D7F7; HTRF with MAB2166 as the donor antibody (**Fig. 8C**) and MSD with MAB2166 as the capture antibody (**Fig. 8D**) paired with MAB5490 or D7F7; HTRF with D7F7 as the donor antibody (**Fig. 8E**) and MSD with D7F7 as the capture antibody (**Fig. 8F**) paired with MAB5490 or MAB2166. The changing pattern of signals with respect to polyQ length was relatively consistent between the HTRF and MSD assays for any given antibody pair, but the inter-sample variability was greater for MSD (**Fig. 8**). None of the assays recapitulated the decrease in mutant HTT levels as detected by western blot; in general, the assays indicated that there was more full-length HTT in heterozygous and/or homozygous *Hdh*Q50 lysates than wild-type, that the levels in the *Hdh*Q80 and *Hdh*Q111 heterozygotes and homozygotes were comparable to wild-type and that decreases in mutant HTT were only apparent in in heterozygous and homozygous CAG140 and zQ175 lysates (**Fig. 8**). However, in all cases, on both platforms, more full-length HTT was detected in the YAC128 lysate than in wild type (**Fig. 8**). Biologically, we do not understand how small changes in the length of the polyQ tract, e.g. from 20 to 50 glutamines, might result in similar changes in the performance of these six HTRF FRET-based and six MSD ELISA-based assays.

**Figure 8.**
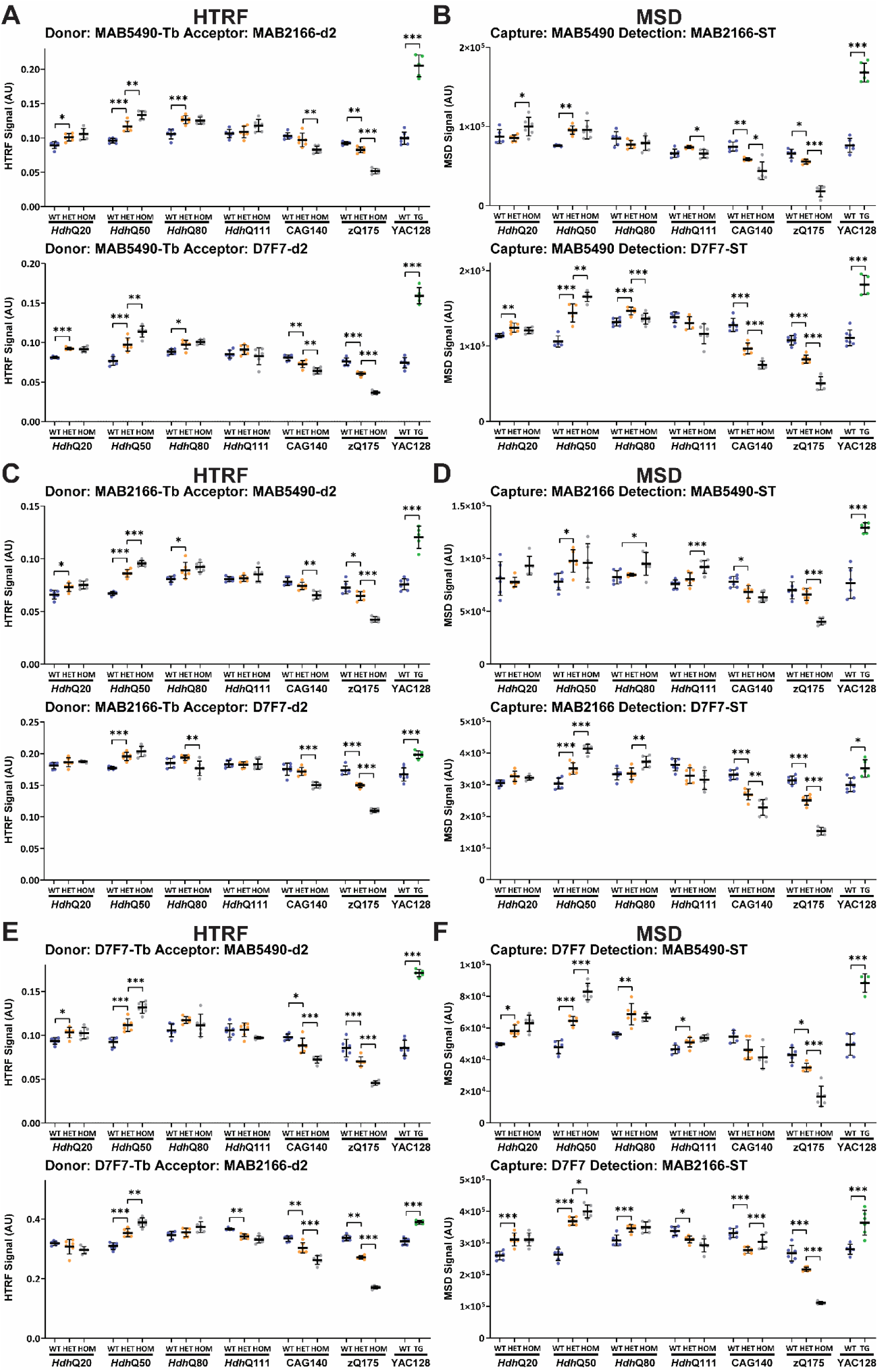
PolyQ-length dependence of HTRF and MSD assays that use pairs of antibodies C-terminal to exon 1 HTT and detect total full-length HTT (mutant and wild-type). Antibody pairings of MAB5490-MAB2166, MAB5490-D7F7 were tested by HTRF **(A)** and MSD **(B)**, of MAB2166-MAB5490 and MAB2166-D7F7 were tested by HTRF **(C)** and MSD **(D)**, and of D7F7-MAB5490 and D7F7-MAB2166 were tested by HTRF **(E)** and MSD **(F)** using cortical lysates from wild-type, heterozygous and homozygous *Hdh*Q20, *Hdh*Q50, *Hdh*Q80, *Hdh*Q111, CAG140 and zQ175 mice at 11 weeks of age and YAC128 and wild-type littermates at 9 weeks of age (n = 6 / genotype). Statistical analysis was Student’s *t*-test or one-way ANOVA with Bonferroni *post hoc* correction per mouse line, mean ± SEM. \**P* ≤ 0.5, \*\**P* ≤ 0.01, \*\*\**P* ≤ 0.001. WT = wild type, HET = heterozygote, HOM = homozygote, TG = transgenic.

## DISCUSSION

Somatic CAG repeat expansion can lead to neurons containing mutant *HTT* with expansions of hundreds of CAGs in the brains of mutation carriers.^6, 7^ The allelic series of Huntington’s disease knock-in mouse models, with repeat lengths ranging from (CAG)_20_ to ∼(CAG)_190_, provide a means by which the molecular and cellular consequences of somatic CAG repeat expansion can be modelled. Here, we have shown that as CAG repeat length increased, the level of cortical full-length mutant HTT protein decreased. Mutant HTT levels were equivalent to those of wild-type HTT in the cortex of mice with (CAG)_50_, decreasing to approximately 10% of wild-type HTT levels with expansions of (CAG)_190_. In contrast, as the CAG repeat length increased, the production of the *Htt1a* transcript and HTT1a protein increased.

We used western blotting to quantify the levels of full-length mutant HTT in cortical tissues from knock-in mice with CAG repeat lengths of approximately 20, 50, 80, 111, 140 and 190 CAGs. We selected the stringent RIPA buffer for lysate preparation, to solubilize membrane bound and hard-to-solubilize proteins. Analysis of heterozygous mice showed that the Huntington’s disease mutation had no effect on wild-type HTT levels. However, whilst the level of mutant HTT in heterozygous mice was equivalent to wild type for *Hdh*Q20 and *Hdh*Q50 mice, this had decreased in the *Hdh*Q80, *Hdh*Q111, CAG140 and zQ175 lines. Cortical mutant HTT levels in homozygous knock-in mice were 50% of wild-type levels in *Hdh*Q111, 23% in CAG140, and only 10% in zQ175 mice. The low HTT levels in homozygous zQ175 mice, are consistent with the recovery of 13% homozygous pups (instead of 25%) from breeding heterozygous male with heterozygous female zQ175 mice (Britt Callahan, personal communication). They are also consistent with data showing that mutant HTT levels are lower than wild type in juvenile Huntington’s disease brains.^36^

This decrease in full-length mutant HTT levels has most likely occurred through a combination of mechanisms. First, our data indicate that the mutant *Htt* transcript was only reduced in the CAG140 and zQ175 cortices, and so this could not have accounted for the much greater reduction in mutant HTT protein observed in the *Hdh*Q80, *Hdh*Q111, CAG140 and zQ175 lines. Second, the translation of the mutant transcript could have been impaired; the translation of transcripts with long CAG repeats has been shown to be decreased due to ribosome stalling and/or collisions.^37^ Third, if soluble mutant HTT had been recruited into HTT aggregates, it would not have been solubilized by RIPA buffer. However, we previously demonstrated that the HTT1a protein is recruited into aggregates in zQ175 brain regions and not full-length HTT.^35^ Fourth, the full-length mutant protein could have undergone proteolysis to generate smaller HTT fragments not detected by D7F7. We have previously assessed the proteolytic fragments generated from mutant HTT in the *Hdh*Q150 knock-in model, but these were not abundant and could not account for such dramatic reductions in mutant HTT levels.^30^

It is essential that soluble and aggregated HTT isoforms can be detected in tissues and biofluids. In our previous study, we had interpreted the detection, by some assays, of much lower levels of mutant HTT in zQ175 striatum and cortex, as compared to *Hdh*Q20, to be due to the expanded polyQ interfering with assay performance.^28^ However, our western blot data suggest that this interpretation was not entirely correct.

In this study, we set out to determine which HTRF and MSD assays might best reflect the changes in mutant HTT levels with increasing CAG repeat length. Cortical tissues from two sets of age-matched knock-in mice from the allelic series were used for all the western blots, HTRF and MSD assays. As mutant HTT can aggregate in tissue lysates, even when stored at -80°C, lysates were always frozen after preparation and used the following day. HTRF and MSD assays were run according to the manufacturer’s recommended conditions. If the results suggested that antibody levels might be limiting, the assay was repeated with a series of lysate dilutions. For a given antibody pair, with a few exceptions, the overall pattern of signals across the allelic series was relatively comparable between HTRF and MSD, although there was often greater sample heterogeneity for MSD.

The validity of the HTRF and MSD assays at detecting full-length HTT was assessed by comparing their performance to our western blot data (**Fig. 1**), for which the protein had been denatured and all antibody binding sites should have been accessible. We used two sets of assays to detect full-length mutant HTT, employing either the MW1 (**Fig. 3**) or 4C9 (**Fig. 4**) antibodies to impart mutant HTT specificity. We would avoid using MW1-based assays because of the non-linear relationship between polyQ-length and signal intensity (**Fig. 3**). The 4C9-based assays better represented the changes in mutant HTT with increasing CAG repeat length; however, none of the assays replicated the western blot data precisely (**Fig. 4**). The choice of assay must be informed by the CAG repeat lengths in the samples that are to be compared. If these are comparable, the choice of assay is less critical. However, if they are known to vary, it is important to know how the assay performance is influenced by CAG repeat length, and the results interpreted with caution.

For the detection of mutant HTT, we would recommend using assays that are selective for either ‘mutant full-length HTT’ or HTT1a and avoid those that detect both isoforms. Comparison of the relative signal levels for the ‘total mutant HTT’ assays (**Fig. 6**) with the patterns obtained for ‘full-length mutant HTT’ (**Figs. 3 and 4**) and for HTT1a (**Fig. 5**) indicated that specific ‘total mutant HTT’ assays preferentially detected one of these isoforms. The most unexpected result was for the frequently used MSD assay, 2B7-MW1, which produced a pattern that tracked with HTT1a; further experiments are needed to confirm this result.

We used two sets of assays to detect ‘total full-length HTT’ levels (wild type and mutant) (**Figs. 7 and 8**). In summary, the HTRF assays that used the 2B7 antibody were the most promising (**Fig. 7**), although none of the 12 HTRF or 12 MSD total full-length HTT assays recapitulate the changes in total HTT with increasing polyQ length as measured by western blot (**Fig. 1**). Instead, we would recommend using species-specific assays i.e. a ‘soluble mutant full length HTT’ assay as discussed above and an ‘endogenous mouse HTT’ assay that we have previously published.^35^

Establishing a set of antibody-based bioassays to detect soluble and aggregated HTT isoforms is complex. These assays are not quantitative, as in the absence of recombinant protein standards with polyQ lengths to match those in the tissue analyte, it is not possible to run standard curves. In order to apply them to any given mouse model or cell line, antibody concentrations and lysate dilutions must be optimized to ensure that an assay is operating in the linear part of the curve.^28^ If performance of an assay can be influenced by polyQ length, it is not possible to compare HTT levels in samples containing HTT with different polyQ repeats. Interpreting the consequences of interventions that might alter polyQ length (e.g. targeting somatic CAG repeat expansion) on HTT levels must be treated with caution as a signal may change because the length of the polyQ repeat has been altered, not because the level of HTT has changed.

This work has significant implications for the development of assays to measure HTT proteins in human biofluids. HTRF and MSD assays are not sufficiently sensitive for use in human plasma or CSF, and instead, SMC assays have been established for that purpose.^21^ Assays that distinguish the mutant HTT isoforms: full-length HTT and HTT1a, as well as assays that distinguish full-length mutant and wild-type HTT are ideally required. In knock-in mice, mutant and wild-type HTT can be distinguished without using antibodies to the polyQ tract, but this is not possible for human samples, and so options are more constrained. Therefore, whether a given assay preferentially selects a specific mutant HTT isoform and / or whether the assay performance is influenced by polyQ-length should be determined. Both questions remain unanswered for the 2B7-MW1 SMC assay that has been frequently used to measure total mutant HTT in CSF,^21, 38^ and could be addressed by applying existing and future assays to diluted tissue lysates from the allelic series of knock-in mice. This is important, because, due to somatic CAG repeat expansion, the mutant HTT protein in CSF may contain polyQ repeats that are heterogeneous in length and may contain up to several hundred glutamine residues.

Our data demonstrate that, as the CAG repeat increases the level of full-length mutant HTT decreases whilst that of the *Htt1a* transcript and the HTT1a protein increase. We have previously shown that the HTT1a is recruited into aggregates,^35^ and that in the striatum of zQ175 mice, further expansion of the CAG repeat is not required for the initiation of HTT aggregation and transcriptional dysregulation.^39^ In heterozygous zQ175 mice, with ∼190 CAGs, cortical full-length mutant HTT is at approximately 10% of wild-type levels, and the extent to which it contributes to the pathogenic process is not known. Single cell analyses from *post-mortem* Huntington’s disease brains have recently shown that somatic expansion occurs in the medium spiny neurons of the striatum^7^ and that expansion of CAG repeats beyond (CAG)_150_ initiates a cell-autonomous toxicity that is executed through the further expansion of the repeat.^40^ Our data predict that levels of full-length mutant HTT in these cells would be very low. If this was the case, therapeutic strategies to lower HTT levels by targeting sequences that are 3’ to exon 1 *HTT* will predominantly decrease wild-type HTT in cells with pathogenic expansions; allele-selective strategies will reduce further what are already very low levels of mutant HTT. Both approaches will leave the pathogenic HTT1a protein levels unchanged.

## ABBREVIATIONS

AlphaLISA: amplified luminescent proximity homogeneous assay
BSA: bovine serum albumin
FELASA: Federation of European Laboratory Animal Science Associations
HDAC4: histone deacetylase 4
HET: heterozygous
HOM: homozygous
HTRF: homogeneous time resolved fluorescence
HTT1a: exon 1 HTT protein
MSD: Meso Scale Discovery
PolyP: polyproline
PolyQ: polyglutamine
Q: glutamine
qPCR: real-time quantitative PCR
SDS- PAGE: SDS-polyacrylamide acrylamide gel electrophoresis
SEM: standard error of the mean
SMC: single molecule counting
WT: wild-type

## ACKNOWLEDGEMENTS

The authors thank Brenda Lager (CHDI Foundation) and Britt Callahan (Jackson Laboratory) for providing mice from the CHDI colonies and Arzo Iqbal for help with mouse dissections. From Revvity, we thank Alexandre Jean for technical discussions and Kerry Chapman for ordering and logistical processing. From MSD, we thank Joao Nunes for technical discussions and Touran Shahbazi for ordering and logistical processing.

## SUPPLEMENTARY MATERIAL

Supplementary material is available at *Brain* online.

## DATA AVAILABILITY

The authors confirm that all the data supporting the findings of this study are available within the article and its supplementary material. Raw data will be shared by the corresponding author on request.

## REFERENCES

1. Bates GP, Dorsey R, Gusella JF, et al. Huntington disease. Nat Rev Dis Primers. Apr 23 2015;1:15005. doi:10.1038/nrdp.2015.5

2. HDCRG. A novel gene containing a trinucleotide repeat that is expanded and unstable on Huntington’s disease chromosomes. The Huntington’s Disease Collaborative Research Group [see comments]. Cell. 1993;72(6):971–983.

3. Duyao M, Ambrose C, Myers R, et al. Trinucleotide repeat length instability and age of onset in Huntington’s disease [see comments]. Nat Genet. 1993;4(4):387–392.

4. Telenius H, Kremer HP, Theilmann J, et al. Molecular analysis of juvenile Huntington disease: the major influence on (CAG)n repeat length is the sex of the affected parent. Hum Mol Genet. 1993;2(10):1535–1540.

5. Telenius H, Kremer B, Goldberg YP, et al. Somatic and gonadal mosaicism of the Huntington disease gene CAG repeat in brain and sperm [published erratum appears in Nat Genet 1994 May;7(1):113]. Nat Genet. 1994;6(4):409–414.

6. Kennedy L, Evans E, Chen CM, et al. Dramatic tissue-specific mutation length increases are an early molecular event in Huntington disease pathogenesis. Hum Mol Genet. Dec 15 2003;12(24):3359–3367.

7. Matlik K, Baffuto M, Kus L, et al. Cell-type-specific CAG repeat expansions and toxicity of mutant Huntingtin in human striatum and cerebellum. Nat Genet. Mar 2024;56(3):383–394. doi:10.1038/s41588-024-01653-6

8. GEM-HD. Identification of Genetic Factors that modify clinical onset of Huntington’s disease. Cell 2015;162:516–526.

9. GeM-HD. CAG Repeat Not Polyglutamine Length Determines Timing of Huntington’s Disease Onset. Cell. Aug 8 2019;178(4):887–900 e14. doi:10.1016/j.cell.2019.06.036

10. Moss DJH, Pardinas AF, Langbehn D, et al. Identification of genetic variants associated with Huntington’s disease progression: a genome-wide association study. Lancet Neurol. Sep 2017;16(9):701–711. doi:10.1016/S1474-4422(17)30161-8

11. Lee JM, Chao MJ, Harold D, et al. A modifier of Huntington’s disease onset at the MLH1 locus. Hum Mol Genet. Oct 1 2017;26(19):3859–3867. doi:10.1093/hmg/ddx286

12. Sathasivam K, Neueder A, Gipson TA, et al. Aberrant splicing of HTT generates the pathogenic exon 1 protein in Huntington disease. Research Support, Non-U.S. Gov’t. Proc Natl Acad Sci U S A. Feb 5 2013;110(6):2366–2370. doi:10.1073/pnas.1221891110

13. Neueder A, Landles C, Ghosh R, et al. The pathogenic exon 1 HTT protein is produced by incomplete splicing in Huntington’s disease patients. Sci Rep. May 2 2017;7(1):1307. doi:10.1038/s41598-017-01510-z

14. Tabrizi SJ, Estevez-Fraga C, van Roon-Mom WMC, et al. Potential disease-modifying therapies for Huntington’s disease: lessons learned and future opportunities. Lancet Neurol. Jul 2022;21(7):645–658. doi:10.1016/S1474-4422(22)00121-1

15. Weiss A, Abramowski D, Bibel M, et al. Single-step detection of mutant huntingtin in animal and human tissues: A bioassay for Huntington’s disease. Anal Biochem. Dec 1 2009;395(1):8–15. doi:S0003-2697(09)00541-7 [pii] 10.1016/j.ab.2009.08.001

16. Baldo B, Paganetti P, Grueninger S, et al. TR-FRET-based duplex immunoassay reveals an inverse correlation of soluble and aggregated mutant huntingtin in huntington’s disease. Chem Biol. Feb 24 2012;19(2):264–275. doi:S1074-5521(12)00016-6 [pii] 10.1016/j.chembiol.2011.12.020

17. Macdonald D, Tessari MA, Boogaard I, et al. Quantification assays for total and polyglutamine-expanded huntingtin proteins. PLoS One. 2014;9(5):e96854. doi:10.1371/journal.pone.0096854

18. Reindl W, Baldo B, Schulz J, et al. Meso scale discovery-based assays for the detection of aggregated huntingtin. PLoS One. 2019;14(3):e0213521. doi:10.1371/journal.pone.0213521

19. Baldo B, Sajjad MU, Cheong RY, et al. Quantification of Total and Mutant Huntingtin Protein Levels in Biospecimens Using a Novel alphaLISA Assay. eNeuro. Jul-Aug 2018;5(4):e2034. doi:10.1523/ENEURO.0234-18.2018

20. Fodale V, Boggio R, Daldin M, et al. Validation of Ultrasensitive Mutant Huntingtin Detection in Human Cerebrospinal Fluid by Single Molecule Counting Immunoassay. J Huntingtons Dis. 2017;6(4):349–361. doi:10.3233/JHD-170269

21. Wild EJ, Boggio R, Langbehn D, et al. Quantification of mutant huntingtin protein in cerebrospinal fluid from Huntington’s disease patients. J Clin Invest. May 2015;125(5):1979–86. doi:10.1172/JCI80743

22. Wheeler VC, Auerbach W, White JK, et al. Length-dependent gametic CAG repeat instability in the Huntington’s disease knock-in mouse. Hum Mol Genet. 1999;8(1):115–122.

23. Menalled LB, Sison JD, Dragatsis I, Zeitlin S, Chesselet MF. Time course of early motor and neuropathological anomalies in a knock-in mouse model of Huntington’s disease with 140 CAG repeats. J Comp Neurol. Oct 6 2003;465(1):11–26.

24. Menalled LB, Kudwa AE, Miller S, et al. Comprehensive Behavioral and Molecular Characterization of a New Knock-In Mouse Model of Huntington’s Disease: zQ175. PLoS One. 2012;7(12):e49838. doi:10.1371/journal.pone.0049838

25. Heikkinen T, Lehtimaki K, Vartiainen N, et al. Characterization of neurophysiological and behavioral changes, MRI brain volumetry and 1H MRS in zQ175 knock-in mouse model of Huntington’s disease. Research Support, Non-U.S. Gov’t. PLoS One. 2012;7(12):e50717. doi:10.1371/journal.pone.0050717

26. Slow EJ, van Raamsdonk J, Rogers D, et al. Selective striatal neuronal loss in a YAC128 mouse model of Huntington disease. Hum Mol Genet. Jul 1 2003;12(13):1555–1567.

27. Fienko S, Landles C, Sathasivam K, et al. Alternative processing of human HTT mRNA with implications for Huntington’s disease therapeutics. Brain. Dec 19 2022;145(12):4409–4424. doi:10.1093/brain/awac241

28. Landles C, Milton RE, Jean A, et al. Development of novel bioassays to detect soluble and aggregated Huntingtin proteins on three technology platforms. Brain Commun. 2021;3(1):fcaa231. doi:10.1093/braincomms/fcaa231

29. Pouladi MA, Stanek LM, Xie Y, et al. Marked differences in neurochemistry and aggregates despite similar behavioural and neuropathological features of Huntington disease in the full-length BACHD and YAC128 mice. Research Support, Non-U.S. Gov’t. Hum Mol Genet. May 15 2012;21(10):2219–2232. doi:10.1093/hmg/dds037

30. Landles C, Sathasivam K, Weiss A, et al. Proteolysis of mutant huntingtin produces an exon 1 fragment that accumulates as an aggregated protein in neuronal nuclei in Huntington disease. J Biol Chem. Mar 19 2010;285(12):8808–8823. doi:M109.075028 [pii] 10.1074/jbc.M109.075028

31. Landles C, Milton RE, Ali N, et al. Subcellular Localization And Formation Of Huntingtin Aggregates Correlates With Symptom Onset And Progression In A Huntington’S Disease Model. Brain Commun. 2020;2(2):fcaa066. doi:10.1093/braincomms/fcaa066

32. Fienko S, Landles C, Sathasivam K, et al. Alternative processing of human HTT mRNA with implications for Huntington’s disease therapeutics. Brain. Jul 6 2022;doi:10.1093/brain/awac241

33. Livak KJ, Schmittgen TD. Analysis of relative gene expression data using real-time quantitative PCR and the 2(-Delta Delta C(T)) Method. Methods. Dec 2001;25(4):402–408. doi:10.1006/meth.2001.1262

34. Langfelder P, Cantle JP, Chatzopoulou D, et al. Integrated genomics and proteomics define huntingtin CAG length-dependent networks in mice. Nat Neurosci. Apr 2016;19(4):623–633. doi:10.1038/nn.4256

35. Smith EJ, Sathasivam K, Landles C, et al. Early detection of exon 1 huntingtin aggregation in zQ175 brains by molecular and histological approaches. Brain Commun. 2023;5(1):fcad010. doi:10.1093/braincomms/fcad010

36. Evers MM, Schut MH, Pepers BA, et al. Making (anti-) sense out of huntingtin levels in Huntington disease. Mol Neurodegener. Apr 28 2015;10:21. doi:10.1186/s13024-015-0018-7

37. Aviner R, Lee TT, Masto VB, Li KH, Andino R, Frydman J. Polyglutamine-mediated ribotoxicity disrupts proteostasis and stress responses in Huntington’s disease. Nat Cell Biol. May 13 2024;doi:10.1038/s41556-024-01414-x

38. Tabrizi SJ, Leavitt BR, Landwehrmeyer GB, et al. Targeting Huntingtin Expression in Patients with Huntington’s Disease. N Engl J Med. May 6 2019;doi:10.1056/NEJMoa1900907

39. Aldous SG, Smith EJ, Landles C, et al. A CAG repeat threshold for therapeutics targeting somatic instability in Huntington’s disease. Brain. Feb 22 2024;doi:10.1093/brain/awae063

40. Handsaker RE, Kashin S, Reed NM, et al. Long somatic DNA-repeat expansion drives neurodegeneration in Huntington’s disease. bioRxiv. 2024;doi:doi.org/10.1101/2024.05.17.592722

