## Supplemental Material for "Mutant HTT protein decreases with CAG repeat expansion: implications for therapeutics and bioassays"

**Supplementary Table 1. CAG repeat sizes of the mice used in this study.**

| Knock-in line | Heterozygotes | Homozygotes<br>(small allele)<br>(large allele) |
|---|---|---|
| *Hdh*Q20 | 18 | 18<br>18 |
| *Hdh*Q50 | 50.1 ± 0.24 | 49.4 ± 0.78<br>49.4 ± 0.78 |
| *Hdh*Q80 | 80.1 ± 0.67 | 80.9 ± 1.51<br>81.4 ± 1.42 |
| *Hdh*Q111 | 114.7 ± 2.38 | 110.4 ± 3.5<br>114.0 ± 2.74 |
| CAG140 | 144.9 ± 2.62 | 140.5 ± 4.85<br>146.8 ± 2.10 |
| zQ175 | 190.3 ± 5.98 | 184.1 ± 5.24<br>194.8 ± 3.96 |

### Supplementary Table 2. Summary of Antibodies.

| Name | Immunogen | Epitope | Species | Reference / Source |
|---|---|---|---|---|
| 2B7 | Human HTT peptide: aa 1-17[1] | LMKAFE* | Mouse Monoclonal | CHDI Foundation |
| MW1 | Human HTT Exon1 (67Q)[2] | PolyQ | Mouse Monoclonal | CHDI Foundation |
| 4C9 | Human HTT peptide: aa 51-71[3] | | Mouse Monoclonal | CHDI Foundation |
| MW8 | AEEPLHRPK (67Q)[2] | Within: AEEPLHRP[4] Ends at proline | Mouse Monoclonal | CHDI Foundation |
| MAB5490 | Human HTT: aa 115-129[5] | Within: QSVRNSPEFQKLLGI (mouse L) | Mouse Monoclonal | Sigma-Aldrich, MAB5490 |
| MAB2166 | HTT fusion protein: aa 181-810[6] | Within aa 443-457 GKVLLGEEEALEDDS [7] | Mouse Monoclonal | Sigma-Aldrich, MAB2166 |
| D7F7 | | Within: aa 1214-1223* QSDTSGPVTT (mouse A) | Rabbit Monoclonal | Cell Signaling Technology #5656 |
| HDAC4 (DM15) | Peptide: human HDAC4 | | Rabbit polyclonal | Merck Life Sciences pAb#H9536 |

*Information provided by the CHDI Foundation

### REFERENCES


1. Weiss A, Abramowski D, Bibel M, et al. Single-step detection of mutant huntingtin in animal and human tissues: A bioassay for Huntington's disease. *Analytical biochemistry*. Dec 1 2009;395(1):8-15. doi:S0003-2697(09)00541-7 [pii]
10.1016/j.ab.2009.08.001
2 .Ko J, Ou S, Patterson PH. New anti-huntingtin monoclonal antibodies: implications for huntingtin conformation and its binding proteins. *Brain research bulletin*. Oct-Nov 1 2001;56(3-4):319-329.
3. Baldo B, Paganetti P, Grueninger S, et al. TR-FRET-based duplex immunoassay reveals an inverse correlation of soluble and aggregated mutant huntingtin in huntington's disease. *Chemistry & biology*. Feb 24 2012;19(2):264-275. doi:S1074-5521(12)00016-6 [pii]
10.1016/j.chembiol.2011.12.020
4. Landles C, Sathasivam K, Weiss A, et al. Proteolysis of mutant huntingtin produces an exon 1 fragment that accumulates as an aggregated protein in neuronal nuclei in Huntington disease. *J Biol Chem*. Mar 19 2010;285(12):8808-8823. doi:M109.075028 [pii]
10.1074/jbc.M109.075028
5. Lunkes A, Lindenberg KS, Ben-Haiem L, et al. Proteases acting on mutant huntingtin generate cleaved products that differentially build up cytoplasmic and nuclear inclusions. *Molecular cell*. Aug 2002;10(2):259-269.
6. Trottier Y, Devys D, Imbert G, et al. Cellular localization of the Huntington's disease protein and discrimination of the normal and mutated form [see comments]. *Nature genetics*. 1995;10(1):104-110.
7. Cong SY, Pepers BA, Roos RA, Van Ommen GJ, Dorsman JC. Epitope mapping of monoclonal antibody 4C8 recognizing the protein huntingtin. *Hybridoma (Larchmt)*. Oct 2005;24(5):231-5. doi:10.1089/hyb.2005.24.231

**Supplementary Table 3. Antibody and lysate concentrations for HTRF assays.**

| Figure Number | Antibody Pairing | Donor (ng / well) | Acceptor (ng / well) | Lysate Dilution Concentration |
|---|---|---|---|---|
| Figure 3 | MW1-Tb : MAB5490-d2 | 1 ng | 20 ng | 5% |
| | MW1-Tb : MAB2166-d2 | 1 ng | 20 ng | 5% |
| | MW1-Tb : D7F7-d2 | 1 ng | 20 ng | 5% |
| | MAB5490-Tb : MW1-d2 | 1 ng | 20 ng | 5% |
| | MAB2166-Tb : MW1-d2 | 1 ng | 20 ng | 5% |
| | D7F7-Tb : MW1-d2 | 1 ng | 20 ng | 5% |
| Figure 4 | 4C9-Tb : MAB5490-d2 | 1 ng | 20 ng | 10% |
| | 4C9-Tb : MAB2166-d2 | 1 ng | 20 ng | 10% |
| | 4C9-Tb : D7F7-d2 | 1 ng | 20 ng | 10% |
| | MAB5490-Tb : 4C9-488 | 1 ng | 20 ng | 10% |
| | MAB2166-Tb : 4C9-488 | 1 ng | 20 ng | 10% |
| | D7F7-Tb : 4C9-488 | 1 ng | 20 ng | 10% |
| Figure 5 | 2B7-Tb : MW8-d2 | 1 ng | 40 ng | 10% |
| | MW1-Tb : MW8-d2 | 1 ng | 40 ng | 10% |
| | 4C9-Tb : MW8-d2 | 1 ng | 20 ng | 10% Crude Lysate |
| Figure 6 | 2B7-Tb : MW1-d2 | 1 ng | 20 ng | 2.5% |
| | 2B7-Tb : 4C9-488 | 1 ng | 20 ng | 5% |
| | MW1-Tb : 2B7-d2 | 1 ng | 20 ng | 2.5% |
| | MW1-Tb : 4C9-488 | 1 ng | 20 ng | 5% |
| | 4C9-Tb : 2B7-d2 | 1 ng | 20 ng | 5% |
| | 4C9-Tb : MW1-d2 | 1 ng | 20 ng | 5% |
| Figure 7 | 2B7-Tb : MAB5490-d2 | 1 ng | 20 ng | 10% |
| | 2B7-Tb : MAB2166-d2 | 1 ng | 20 ng | 10% |
| | 2B7-Tb : D7F7-d2 | 1 ng | 20 ng | 10% |
| | MAB5490-Tb : 2B7-d2 | 1 ng | 20 ng | 10% |
| | MAB2166-Tb : 2B7-d2 | 1 ng | 20 ng | 10% |
| | D7F7-Tb : 2B7-d2 | 1 ng | 20 ng | 10% |
| Figure 8 | MAB5490-Tb : MAB2166-d2 | 1 ng | 20 ng | 5% |
| | MAB5490-Tb : D7F7-d2 | 1 ng | 20 ng | 5% |
| | MAB2166-Tb : MAB5490-d2 | 1 ng | 20 ng | 5% |
| | MAB2166-Tb : D7F7-d2 | 1 ng | 20 ng | 5% |
| | D7F7-Tb : MAB5490-d2 | 1 ng | 20 ng | 5% |
| | D7F7-Tb : MAB2166-d2 | 1 ng | 20 ng | 5% |

NB: Final antibody working conditions and lysate dilutions must always be established by end user, since plate reader sensitivities will change depending on make/model.

**Supplementary Table 4. Antibody and lysate concentrations for MSD assays.**

| Figure Number | Antibody Pairing | Capture | Sulfo-tag (μg/mL) | Lysate Dilution Concentration |
|---|---|---|---|---|
| Figure 3 | MW1-Capture : MAB5490-ST | MSD default | 2.5 μg/mL | 12% |
| | MW1-Capture : MAB2166-ST | MSD default | 2.5 μg/mL | 12% |
| | MW1-Capture : D7F7-St | MSD default | 2.5 μg/mL | 12% |
| | MAB5490-Capture : MW1-ST | MSD default | 1.5 μg/mL | 12% |
| | MAB2166-Capture : MW1-ST | MSD default | 1.5 μg/mL | 12% |
| | D7F7-Capture : MW1-ST | MSD default | 1.5 μg/mL | 12% |
| Figure 4 | 4C9-Capture : MAB5490-ST | MSD default | 2.5 μg/mL | 12% |
| | 4C9-Capture : MAB2166-ST | MSD default | 2.5 μg/mL | 12% |
| | 4C9-Capture : D7F7-ST | MSD default | 2.5 μg/mL | 12% |
| | MAB5490-Capture : 4C9-ST | MSD default | 2.5 μg/mL | 12% |
| | MAB2166-Capture : 4C9-ST | MSD default | 2.5 μg/mL | 12% |
| | D7F7-Capture : 4C9-ST | MSD default | 2.5 μg/mL | 12% |
| Figure 5 | 2B7-Capture : MW8-ST | MSD default | 2.5 μg/mL | 18% |
| | MW1-Capture : MW8-ST | MSD default | 2.5 μg/mL | 18% |
| | 4C9-Capture : MW8-ST | MSD default | 2.5 μg/mL | 18% Crude Lysate |
| Figure 6 | 2B7-Capture : MW1-ST | MSD default | 1.5 μg/mL | 12% |
| | 2B7-Capture : 4C9-ST | MSD default | 2.5 μg/mL | 12% |
| | MW1-Capture : 2B7-ST | MSD default | 2.5 μg/mL | 12% |
| | MW1-Capture : 4C9-ST | MSD default | 2.5 μg/mL | 12% |
| | 4C9-Capture : 2B7-ST | MSD default | 2.5 μg/mL | 12% |
| | 4C9-Capture : MW1-ST | MSD default | 1.5 μg/mL | 12% |
| Figure 7 | 2B7-Capture : MAB5490-ST | MSD default | 2.5 μg/mL | 12% |
| | 2B7-Capture : MAB2166-ST | MSD default | 2.5 μg/mL | 12% |
| | 2B7-Capture : D7F7-ST | MSD default | 2.5 μg/mL | 12% |
| | MAB5490-Capture : 2B7-ST | MSD default | 2.5 μg/mL | 12% |
| | MAB2166-Capture : 2B7-ST | MSD default | 2.5 μg/mL | 12% |
| | D7F7-Capture : 2B7-ST | MSD default | 2.5 μg/mL | 12% |
| Figure 8 | MAB5490-Capture : MAB2166-ST | MSD default | 2.5 μg/mL | 12% |
| | MAB5490-Capture: D7F7-ST | MSD default | 2.5 μg/mL | 12% |
| | MAB2166-Capture : MAB5490-ST | MSD default | 2.5 μg/mL | 12% |
| | MAB2166-Capture : D7F7-ST | MSD default | 2.5 μg/mL | 12% |
| | D7F7-Capture : MAB5490-ST | MSD default | 2.5 μg/mL | 12% |
| | D7F7-Capture : MAB2166-ST | MSD default | 2.5 μg/mL | 12% |

**Supplementary Table 5. PCR primers and probes.**

| Assay | Forward primer | Reverse primer | Probe |
|---|---|---|---|
| *FL HTT* (3'UTR) | CCT GGT ATG TGG ATC AGA AGT C | CAG GTA TGC CTA CTG GGT AAA T | 5' FAM -AGC TCT TGC CAG ATG GTT CTG AGC 3' BHQ1 |
| *HTT1a* | TCCTCATCAGGCCTAAGAGC TGG | GAGACCTCCTAAAAGC ATTATGTCATC | 5' FAM AGTGCAGGACA GCGTGA 3' BHQ1 |

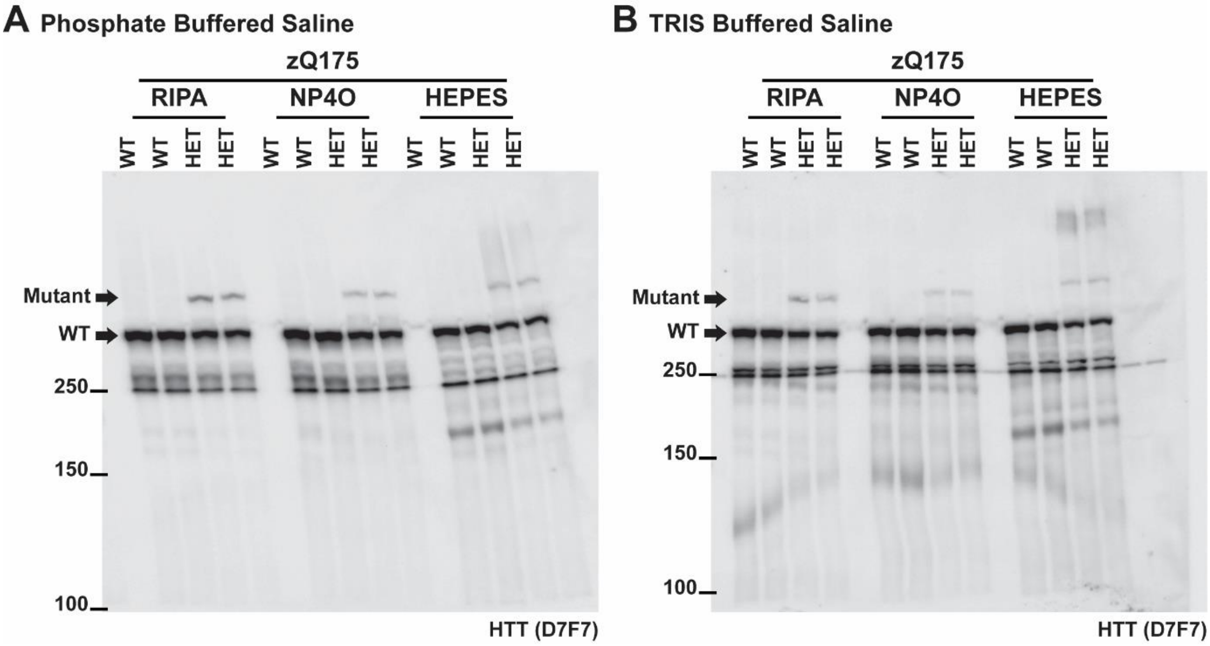


**Supplementary Figure 1. Comparison of lysate buffers on the immunodetection of HTT with the D7F7 antibody on western blots of cortical lysates.** Cortical lysates from wild-type and heterozygous zQ175 mice (n = 2 / genotype) prepared in RIPA, NP40 or HEPES buffers, separated by 7.5% SDS-PAGE, and immunoprobed with D7F7 in **(A)** PBST (phosphate buffered saline, 0.1% TWEEN 20) or **(B)** TBST (TRIS-buffered saline, 0.1% TWEEN 20). Blots immunoprobed in PBST were cleaner than those probed in TBST. The mutant HTT signal was marginally greater for lysates prepared in RIPA buffer and there was less non-specific background for the lysates prepared in RIPA buffer. WT = wild type, HET = heterozygote, size markers are in kDa.

### Supplementary Figure 2

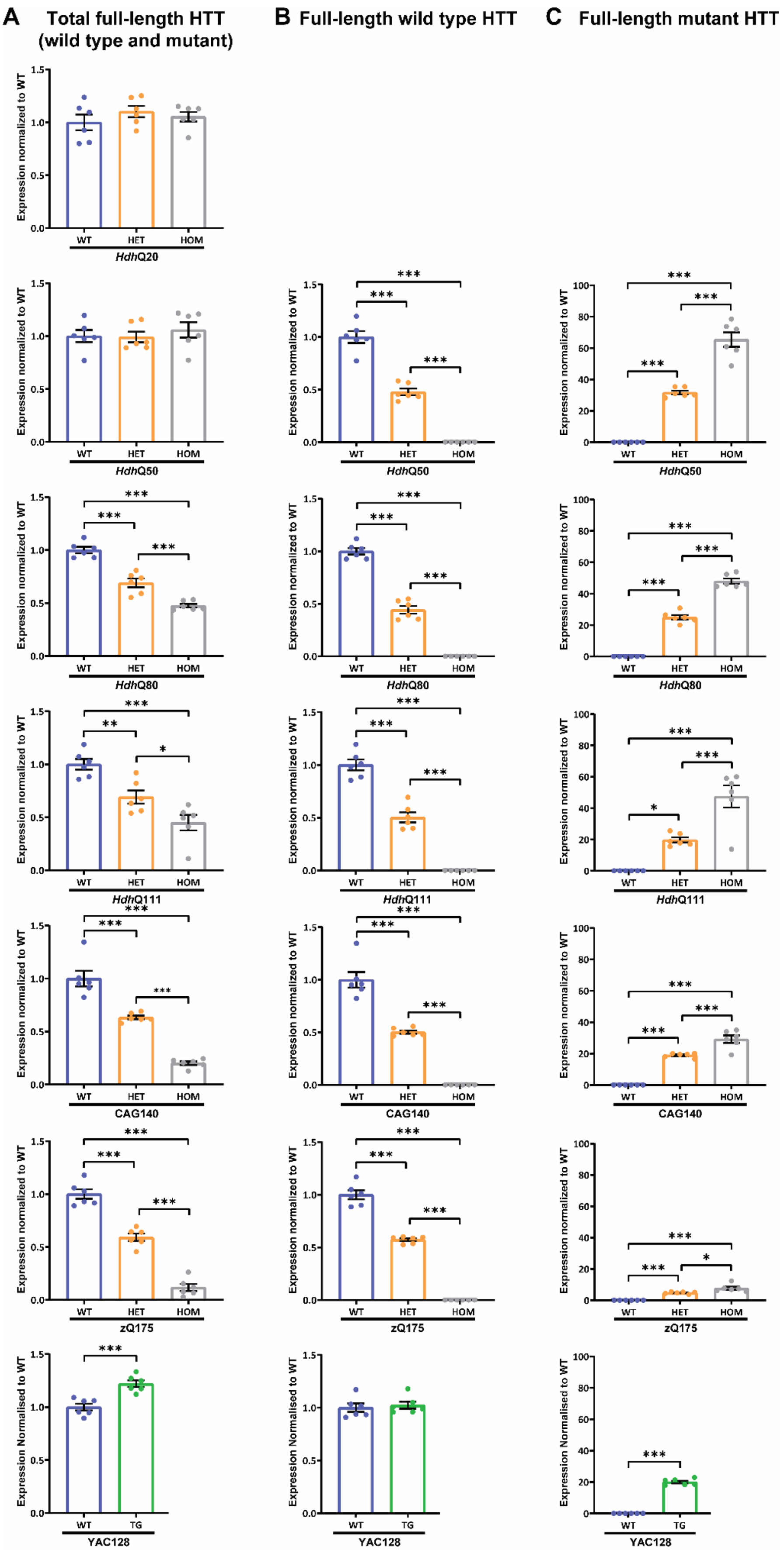

**Supplementary Figure 2. Quantification of full-length wild-type and mutant HTT in cortical lysates in the allelic series of knock-in mouse lines and YAC128 mice.** Quantification of the western blots in **Fig. 1 and Supplementary Figs 4–10**. **(A)** The levels of mutant and wild-type HTT were equivalent in the *Hdh*Q20 and *Hdh*Q50 lines. In contrast, the level of mutant HTT in the *Hdh*Q80, *Hdh*Q111, CAG140 and zQ175 cortical lysates decreased with increasing CAG repeat length. There was approximately 20% more HTT in YAC128 mice as compared to wild type. **(B)** Quantification of the wild-type HTT signals in the heterozygous lysates confirmed that these were approximately 50% of that in wild-type mice. Wild-type HTT was at endogenous levels in YAC128 mice. **(C)** Quantification of the mutant HTT signals in the homozygotes confirmed that these were approximately double that detected in the heterozygous lysates for all lines. Statistical analysis was by Student's *t*-test and one-way ANOVA with Bonferroni *post hoc* correction. Error bars are mean ± SEM. $*P \leq 0.05$, $**P \leq 0.01$, $***P \leq 0.001$. WT = wild type, HET = heterozygote, HOM = homozygote, TG = transgenic.

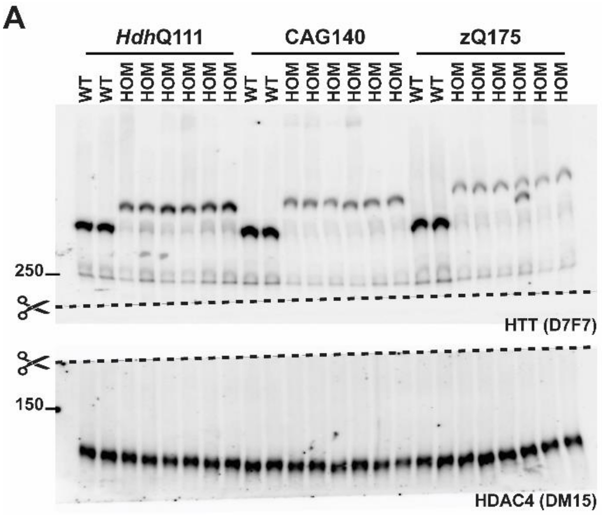


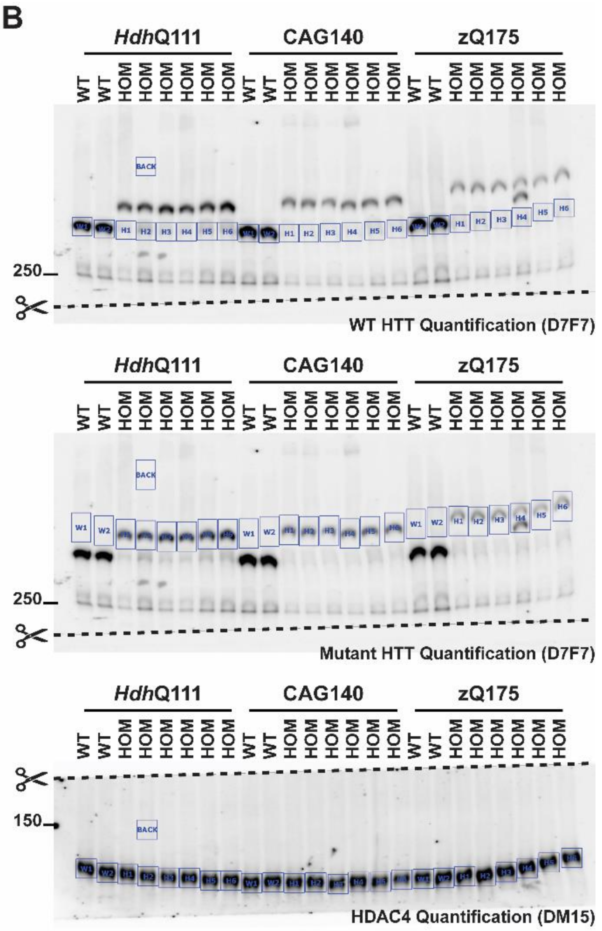


D

| Mouse knock-in line | Total full-length HTT (WT or mutant), normalized to WT | |
|---|---|---|
| | WT | HOM |
| *Hdh*Q111 | 100 | 50 |
| CAG140 | 100 | 23 |
| zQ175 | 100 | 10 |

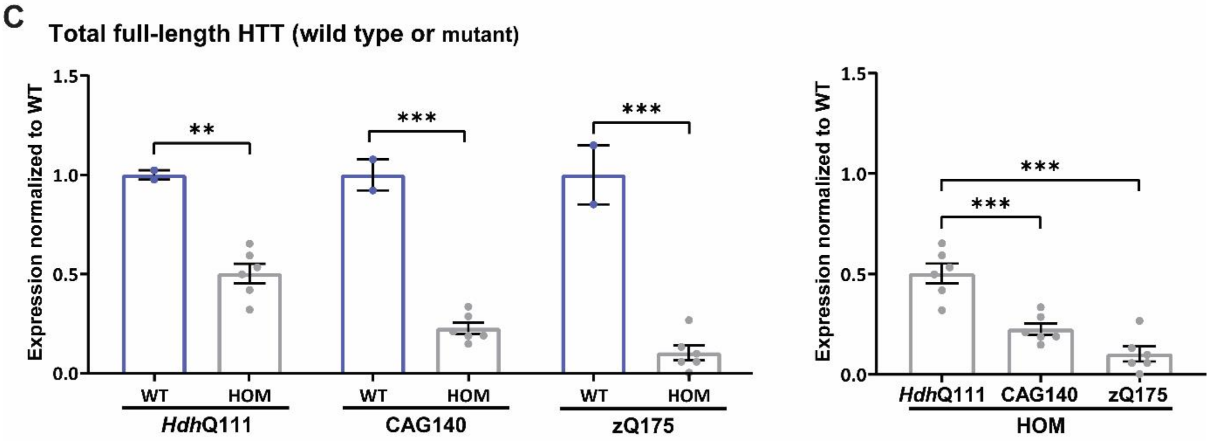


**Supplementary Figure 3. Comparison of mutant HTT levels in the *Hdh*Q111, CAG140 and zQ175 cortical lysates. (A)** The western blot was cut where indicated. The top section probed with D7F7 for HTT, and the bottom section with DM15 for HDAC4 (n = 6/genotype). **(B)** The signals were developed using a ChemiDoc-MP Imaging System (Bio-Rad) and quantified using Image Lab Software (Bio-Rad) as illustrated. **(C)** Mutant HTT levels in the homozygous lysates were normalized to wild-type HTT levels. Statistical analysis was by Student's *t*-test and one-way ANOVA with Bonferroni *post hoc* correction. Error bars are mean ± SEM. $**P \leq 0.01$, $***P \leq 0.001$. WT = wild type, HOM = homozygote, BACK = background. Protein size markers are in kDa.

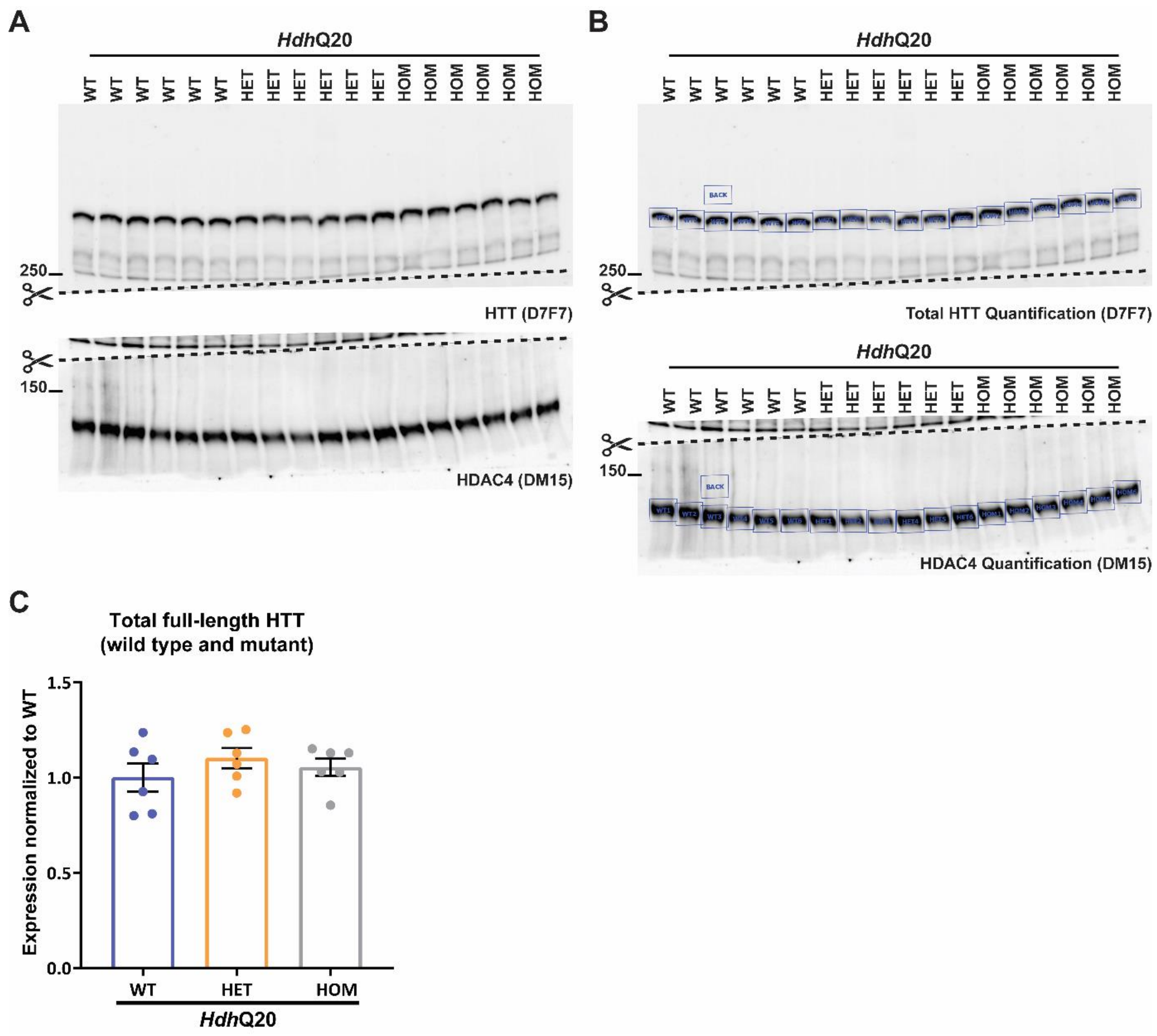


**Supplementary Figure 4. Quantification of full-length HTT in cortical lysates from wild-type, heterozygous and homozygous mice from the *Hdh*Q20 colony. (A)** The western blot was cut where indicated. The top section was probed with D7F7 for HTT, and the bottom section with DM15 for HDAC4 (n = 6/genotype). **(B)** The signals were developed using a ChemiDoc-MP Imaging System (Bio-Rad) and quantified using Image Lab Software (Bio-Rad) as illustrated. **(C)** Quantitation of total full-length HTT in the heterozygous and homozygous lysates normalized to wild-type levels. WT = wild type, HET = heterozygote, HOM = homozygote, BACK = background. Protein size markers are in kDa.

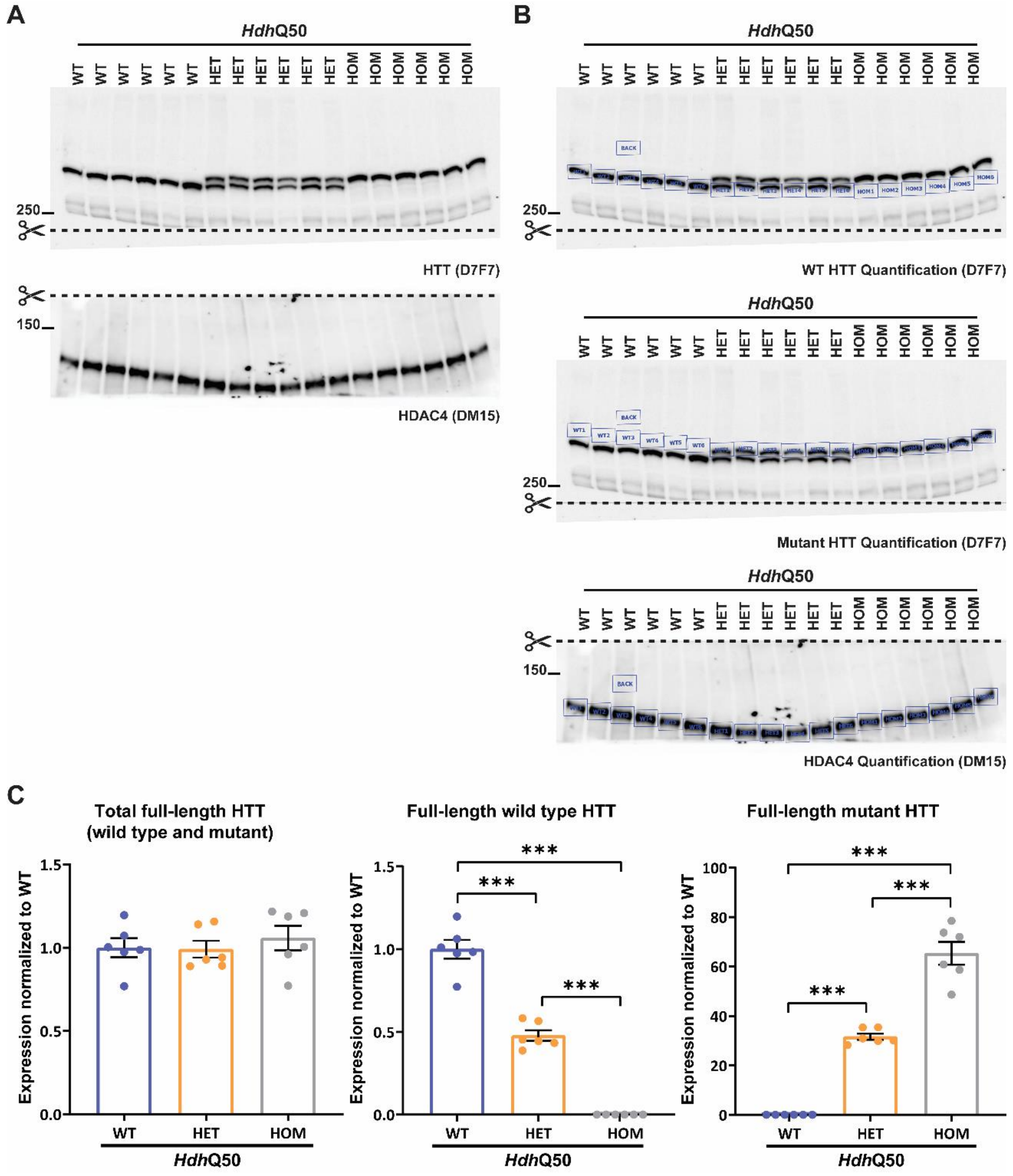


**Supplementary Figure 5. Quantification of full-length HTT in cortical lysates from wild-type, heterozygous and homozygous mice from the *Hdh*Q50 colony. (A)** The western blot was cut where indicated. The top section probed with D7F7 for HTT, and the bottom section with DM15 for HDAC4 (n = 6/genotype). **(B)** The signals were developed using a ChemiDoc-MP Imaging System (Bio-Rad) and quantified using Image Lab Software (Bio-Rad) as illustrated. **(C)** Total full-length HTT and full-length wild-type HTT in the heterozygous and homozygous lysates were normalized to wild-type HTT levels. Full-length mutant HTT levels in the heterozygous and homozygous lysates were normalized to wild-type background. Statistical analysis was by one-way ANOVA with Bonferroni *post hoc* correction. Error bars are mean ± SEM. ****P* ≤ 0.001. WT = wild type, HET = heterozygote, HOM = homozygote, BACK = background. Protein size markers are in kDa.

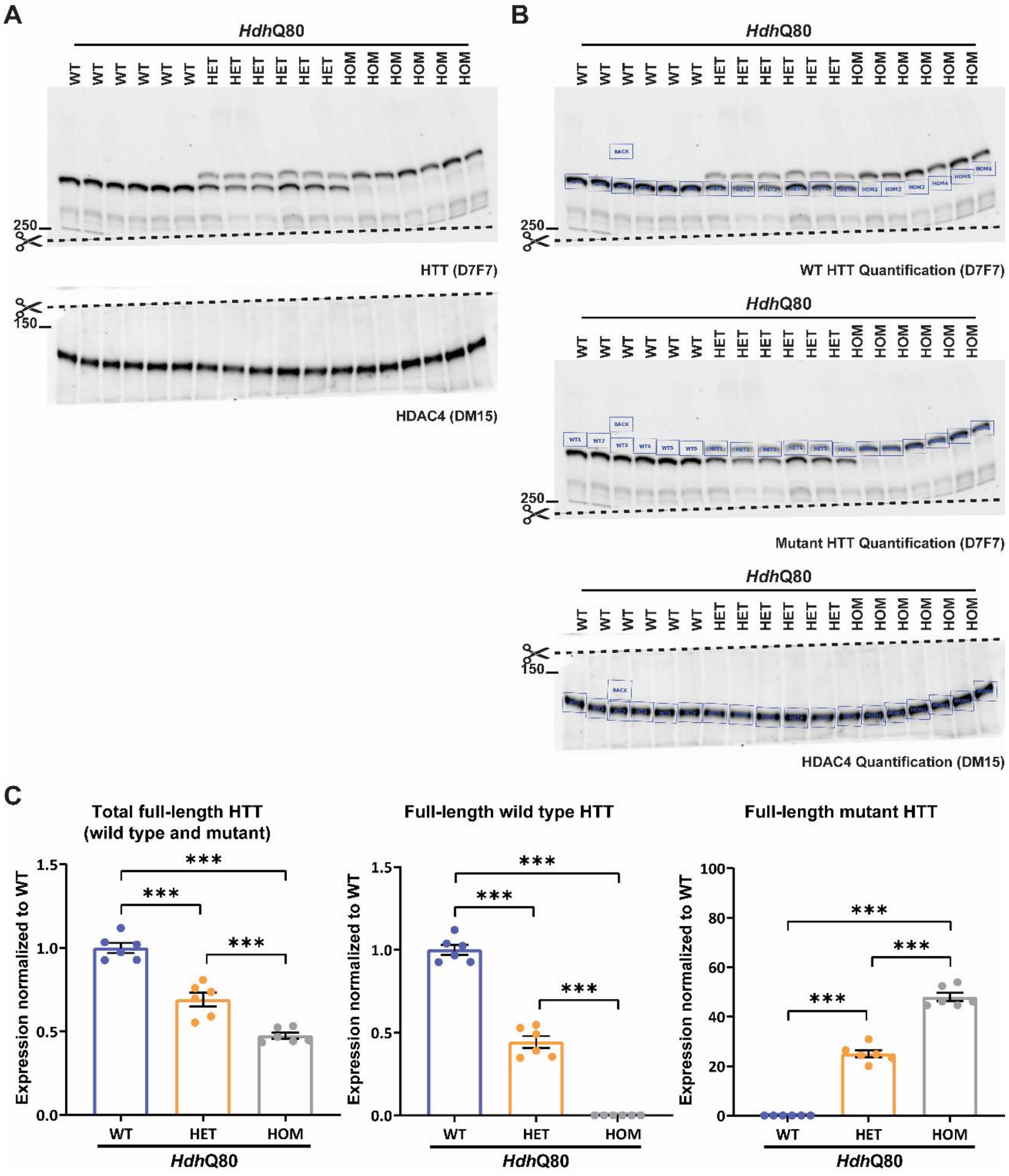


**Supplementary Figure 6. Quantification of full-length HTT in cortical lysates from wild-type, heterozygous and homozygous mice from the *Hdh*Q80 colony. (A)** The western blot was cut where indicated. The top section probed with D7F7 for HTT, and the bottom section with DM15 for HDAC4 (n = 6/genotype). **(B)** The signals were developed using a ChemiDoc-MP Imaging System (Bio-Rad) and quantified using Image Lab Software (Bio-Rad) as illustrated. **(C)** Total full-length HTT and full-length wild-type HTT in the heterozygous and homozygous lysates were normalized to wild-type HTT levels. Full-length mutant HTT levels in the heterozygous and homozygous lysates were normalized to wild-type background. Statistical analysis was by one-way ANOVA with Bonferroni *post hoc* correction. Error bars are mean ± SEM. ***$P \leq 0.001$. WT = wild type, HET = heterozygote, HOM = homozygote, BACK = background. Protein size markers are in kDa.

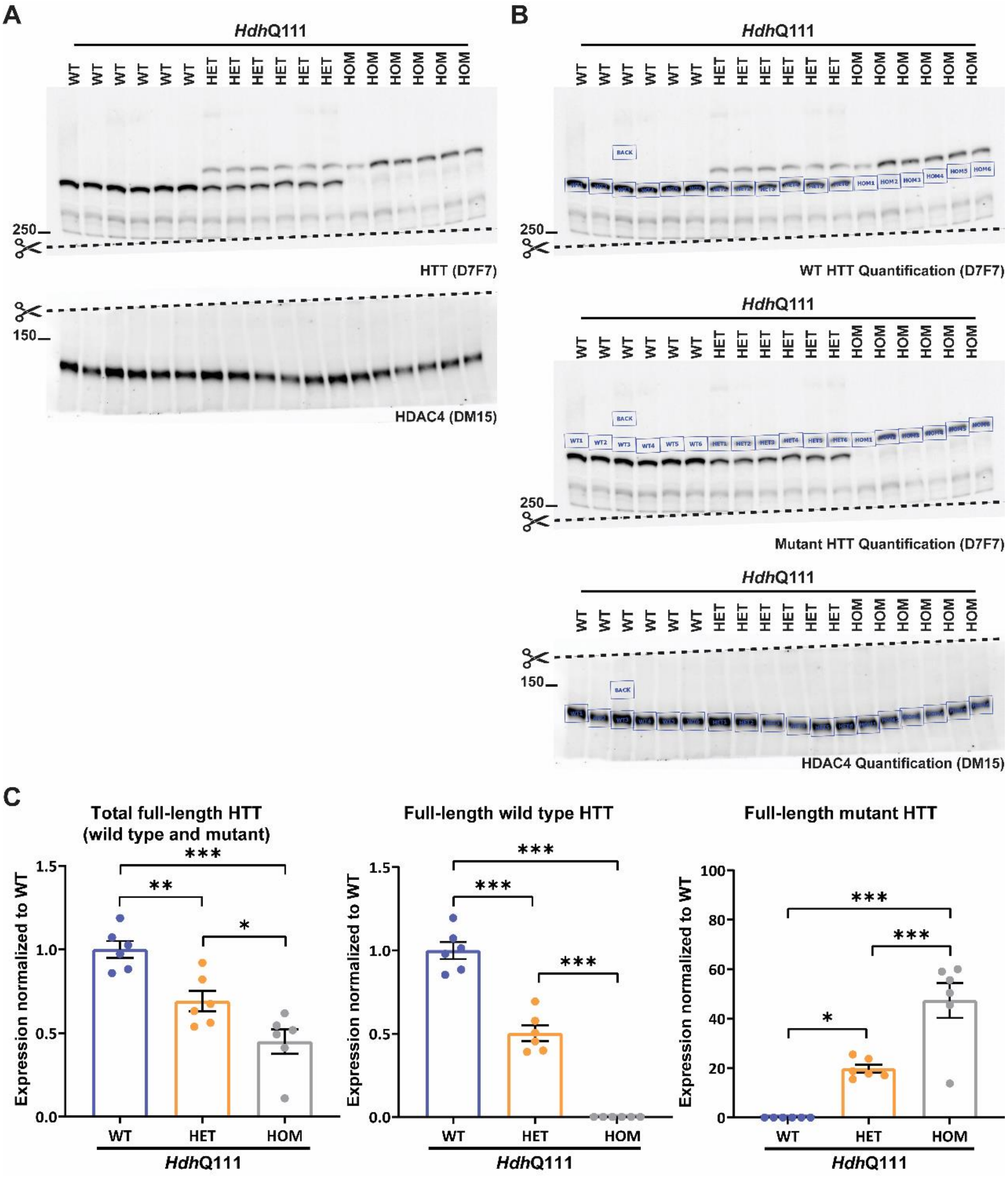


**Supplementary Figure 7. Quantification of full-length HTT in cortical lysates from wild-type, heterozygous and homozygous mice from the *Hdh*Q111 colony. (A)** The western blot was cut where indicated. The top section probed with D7F7 for HTT, and the bottom section with DM15 for HDAC4(n = 6/genotype). **(B)** The signals were developed using a ChemiDoc-MP Imaging System (Bio-Rad) and quantified using Image Lab Software (Bio-Rad) as illustrated. **(C)** Total full-length HTT and full-length wild-type HTT in the heterozygous and homozygous lysates were normalized to wild-type HTT levels. Full-length mutant HTT levels in the heterozygous and homozygous lysates were normalized to wild-type background. Statistical analysis was by one-way ANOVA with Bonferroni *post hoc* correction. Error bars are mean ± SEM. $^{*}P \leq 0.05$, $^{**}P \leq 0.01$, $^{***}P \leq 0.001$. WT = wild type, HET = heterozygote, HOM = homozygote, BACK = background. Protein size markers are in kDa.

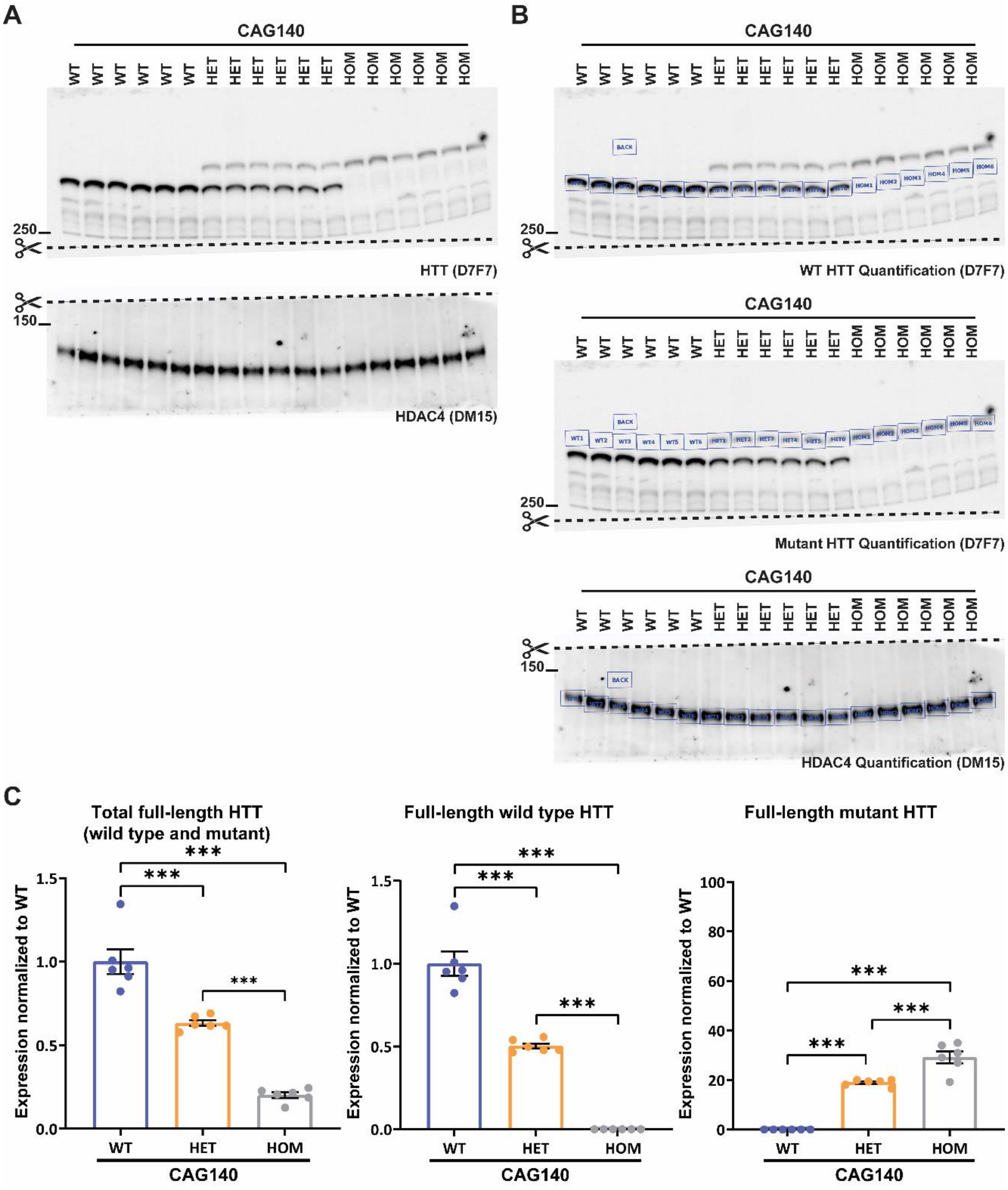


**Supplementary Figure 8. Quantification of full-length HTT in cortical lysates from wild-type, heterozygous and homozygous mice from the CAG140 colony. (A)** The western blot was cut where indicated. The top section probed with D7F7 for HTT, and the bottom section with DM15 for HDAC4 (n = 6/genotype). **(B)** The signals were developed using a ChemiDoc-MP Imaging System (Bio-Rad) and quantified using Image Lab Software (Bio-Rad) as illustrated. **(C)** Total full-length HTT and full-length wild-type HTT in the heterozygous and homozygous lysates were normalized to wild-type HTT levels. Full-length mutant HTT levels in the heterozygous and homozygous lysates were normalized to wild-type background. Statistical analysis was by one-way ANOVA with Bonferroni *post hoc* correction. Error bars are mean ± SEM. ****P* ≤ 0.001. WT = wild type, HET = heterozygote, HOM = homozygote, BACK = background. Protein size markers are in kDa.

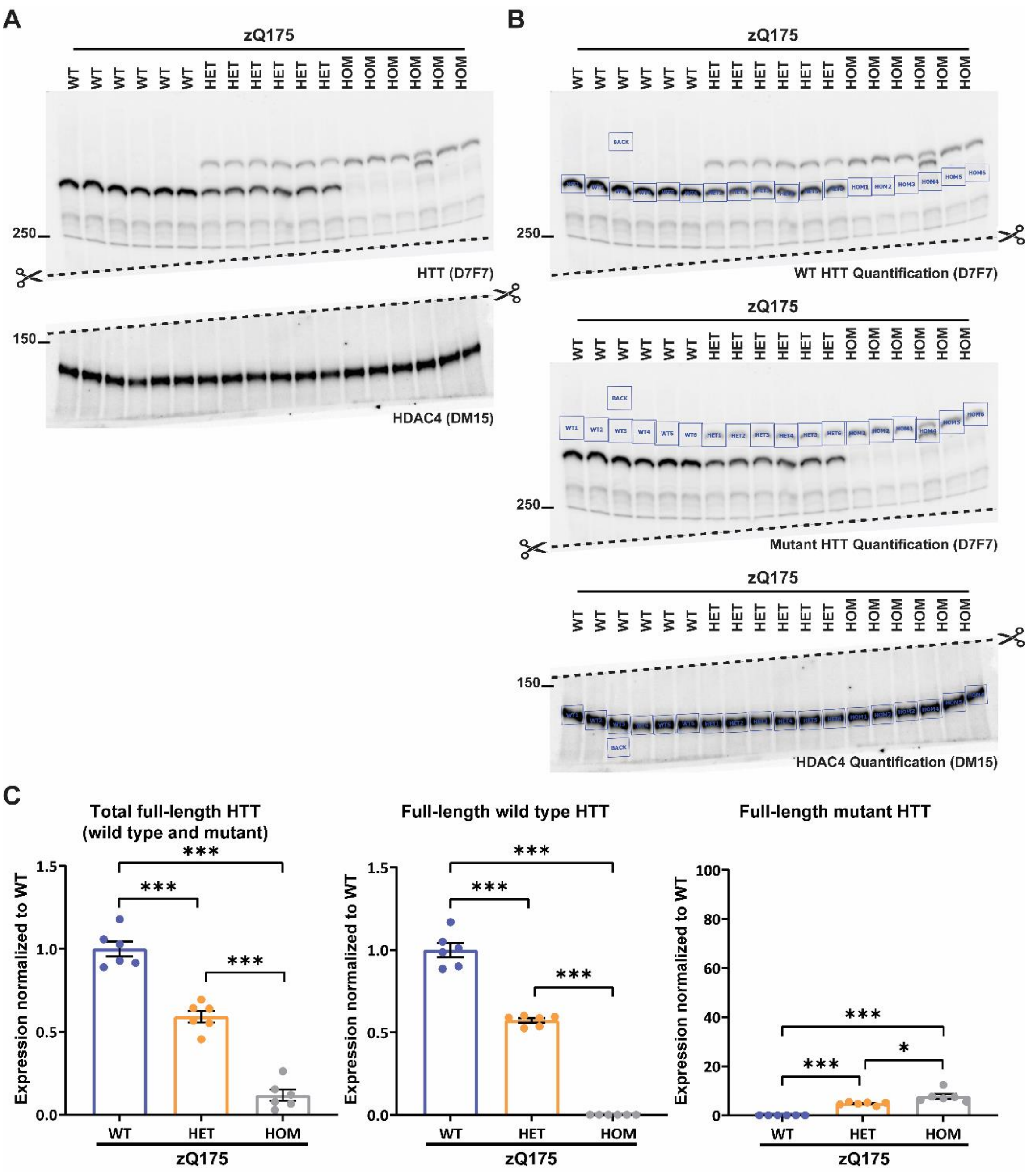


**Supplementary Figure 9. Quantification of full-length HTT in cortical lysates from wild-type, heterozygous and homozygous mice from the zQ175 colony. (A)** The western blot was cut where indicated. The top section probed with D7F7 for HTT, and the bottom section with DM15 for HDAC4 (n = 6/genotype). **(B)** The signals were developed using a ChemiDoc-MP Imaging System (Bio-Rad) and quantified using Image Lab Software (Bio-Rad) as illustrated. **(C)** Total full-length HTT and full-length wild-type HTT in the heterozygous and homozygous lysates were normalized to wild-type HTT levels. Full-length mutant HTT levels in the heterozygous and homozygous lysates were normalized to wild-type background. Statistical analysis was by one-way ANOVA with Bonferroni *post hoc* correction. Error bars are mean ± SEM. **P* ≤ 0.05, ****P* ≤ 0.001. WT = wild type, HET = heterozygote, HOM = homozygote, BACK = background. Protein size markers are in kDa.

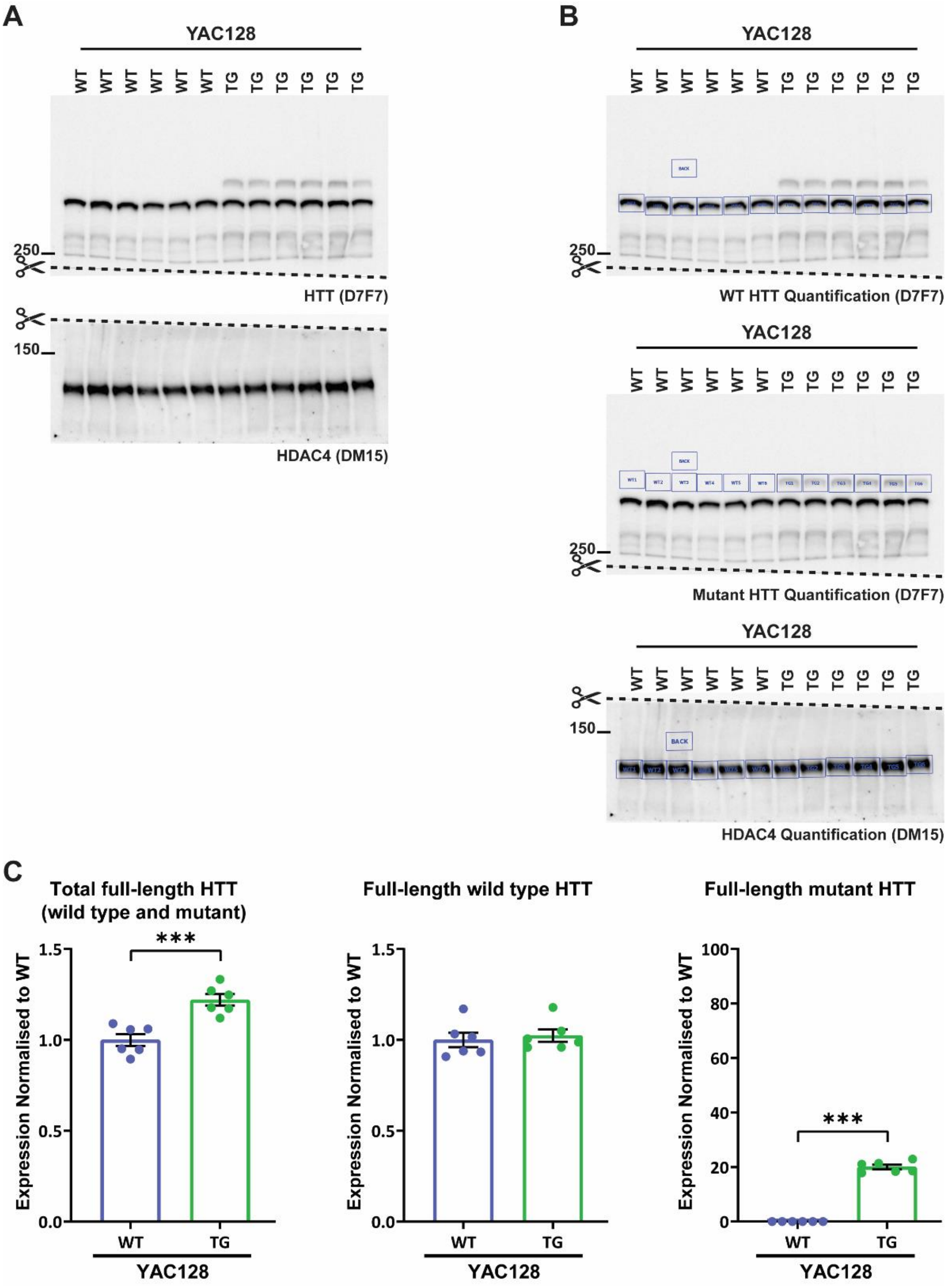


**Supplementary Figure 10. Quantification of full-length HTT in cortical lysates from wild-type and transgenic mice from the YAC128 colony.** **(A)** The western blot was cut where indicated. The top section probed with D7F7 for HTT, and the bottom section with DM15 for HDAC4 (n = 6/genotype). **(B)** The signals were developed using a ChemiDoc-MP Imaging System (Bio-Rad) and quantified using Image Lab Software (Bio-Rad) as illustrated. **(C)** Total full-length HTT and full-length wild-type HTT in the transgenic lysates were normalized to wild-type HTT levels. Full-length mutant HTT levels in the transgenic lysates were normalized to wild-type background. Statistical analysis was by Student's *t*-test. Error bars are mean ± SEM. $***P \leq 0.001$. WT = wild type, TG = transgenic, BACK = background. Protein size markers are in kDa.

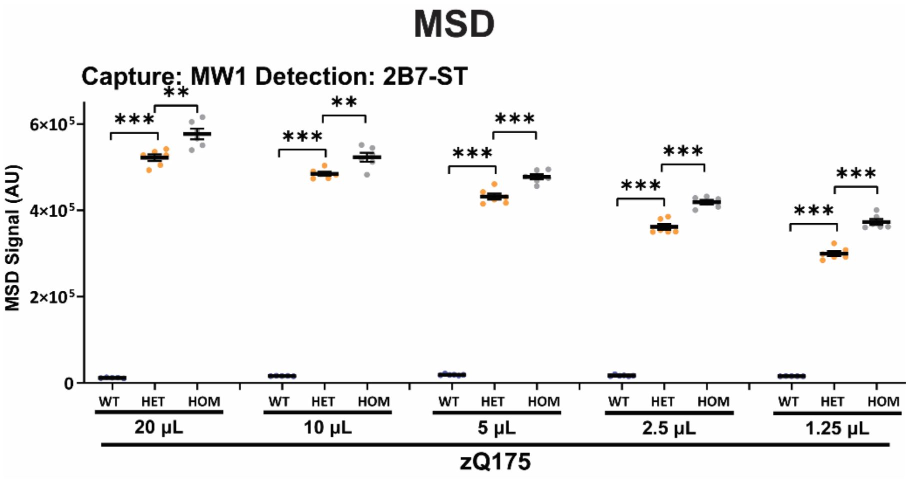


**Supplementary Figure 11. The MW1-2B7 MSD signal was not saturated in lysates from heterozygous knock-in mice.** The antibody pairing of MW1-2B7 was tested by MSD in decreasing serial dilutions of cortical lysates from wild-type, heterozygous and homozygous zQ175 mice at 11 weeks of age (n = 6 / genotype). The signal was always higher in the homozygous lysates, indicating that the heterozygous signal was not saturated. Statistical analysis was Student's *t*-test per lysate dilution, mean ± SEM. **$P \leq 0.01$, ***$P \leq 0.001$. WT = wild type, HET = heterozygote, HOM = homozygote.

### Supplementary Figure 12.

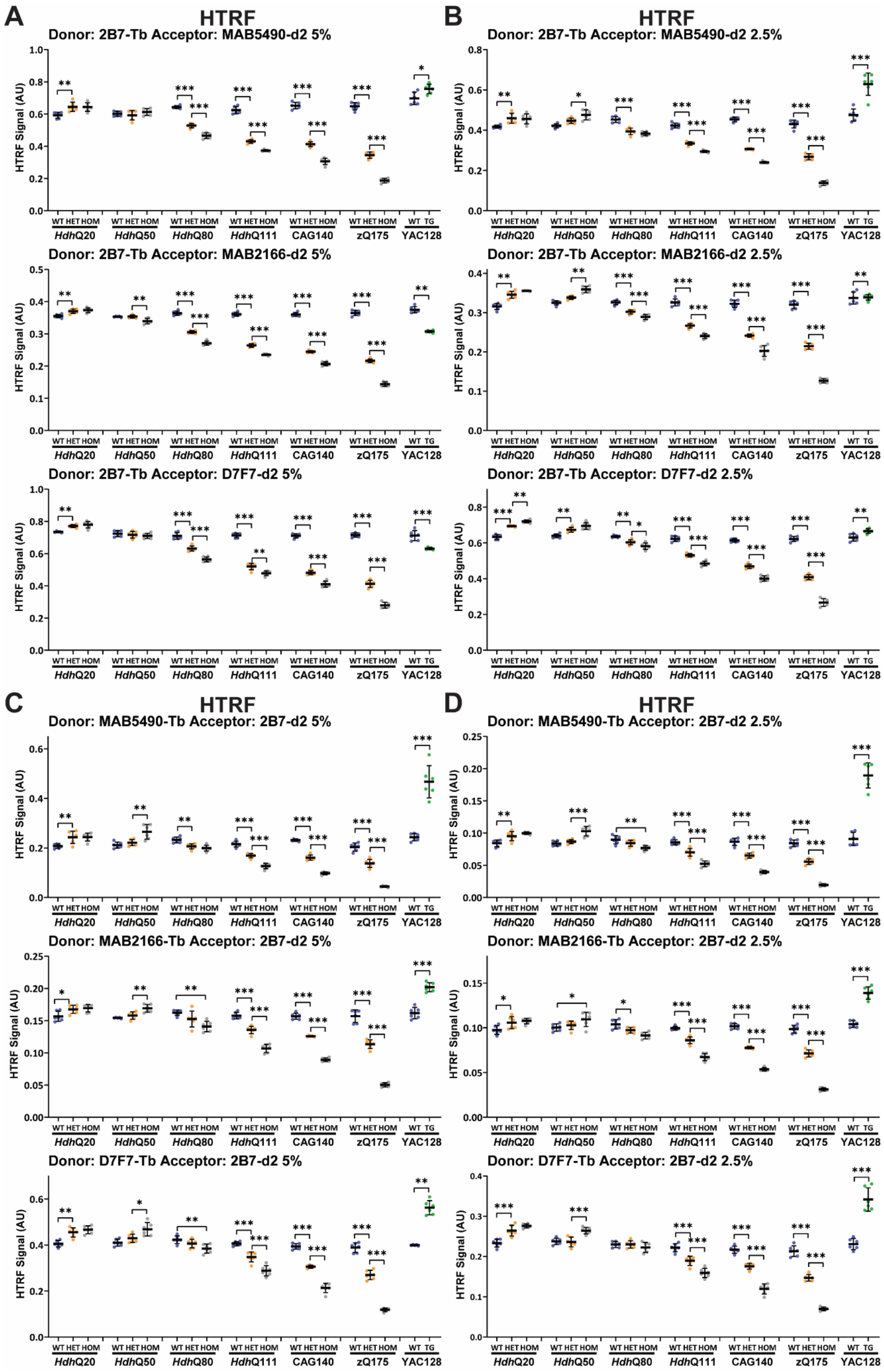

**Supplementary Figure 12. The effect of lysate concentration on the HTRF assays that use 2B7 and detect total full-length HTT (mutant and wild type).** Antibody pairings of 2B7-MAB5490, 2B7-MAB2166 and 2B7-D7F79 were tested by HTRF with 5% lysate **(A)** and 2.5% lysate **(B),** and of MAB5490-2B7, MAB2166-2B7 and D7F7-2B7 were tested by HTRF with 5% lysate **(C)** and 2.5% lysate **(D)** using cortical lysates from wild-type, heterozygous and homozygous *Hdh*Q20, *Hdh*Q50, *Hdh*Q80, *Hdh*Q111, CAG140 and zQ175 mice at 11 weeks of age and YAC128 and wild-type littermates at 9 weeks of age (n = 6 / genotype). Statistical analysis was Student's *t*-test or one-way ANOVA with Bonferroni *post hoc* correction per mouse line, mean ± SEM. *$P \leq 0.5$, **$P \leq 0.01$, ***$P \leq 0.001$. WT = wild type, HET = heterozygote, HOM = homozygote, TG = transgenic.
